# Modeling compound lipid homeostasis using stable isotope tracing

**DOI:** 10.1101/2024.10.16.618599

**Authors:** Karl Wessendorf-Rodriguez, Maureen L. Ruchhoeft, Ethan L. Ashley, Hector M. Galvez, Christopher W. Murray, Yale Huang, Grace H. McGregor, Shrikaar Kambhampati, Reuben J. Shaw, Christian M. Metallo

## Abstract

Lipids represent the most diverse pool of metabolites found in cells, facilitating compartmentation, signaling, and other functions. Dysregulation of lipid metabolism is linked to disease states such as cancer and neurodegeneration. However, limited tools are available for quantifying metabolic fluxes across the lipidome. To directly measure reaction fluxes encompassing compound lipid homeostasis, we applied stable isotope tracing, liquid chromatography-high-resolution mass spectrometry, and network-based isotopologue modeling to non-small cell lung cancer (NSCLC) models. Compound lipid metabolic flux analysis (CL-MFA) enables the concurrent quantitation of fatty acid synthesis, elongation, headgroup assembly, and salvage reactions within virtually any biological system. Here, we resolve liver kinase B1 (LKB1)-mediated regulation of sphingolipid recycling in NSCLC cells and precision-cut lung slice cultures. We also demonstrate that widely used tissue culture conditions drive cells to upregulate fatty acid synthase flux to supraphysiological levels. Finally, we identify previously uncharacterized isozyme specificity of ceramide synthase inhibitors. These results highlight the ability of CL-MFA to quantify lipid cycling in biological systems to discover biological function and elucidate molecular mechanisms in membrane lipid metabolism.

## Introduction

Compound lipids are the most abundant and diverse group of metabolites in cells and tissues. They facilitate numerous biological processes that include membrane compartmentalization, bioenergetics, signal transduction, protein function, and cell-cell interactions^1–6^. Whereas sterols exhibit more limited biochemical modifications, compound lipids like glycerolipids and sphingolipids are highly metabolized and processed in pathways such as the Lands cycle^7^. They are generally synthesized from a head group and acyl-chains that are further modified and catabolized by diverse enzymes spanning numerous organelles. The physical properties and function of each lipid therefore depend on the backbone, head group chemistry, and acyl-chain structure (chain length, desaturation, hydroxylation). For example, the incorporation of long chain versus very-long chain fatty acids (VLCFAs) into ceramides (Cer) impacts their trafficking, fate and function^8–10^. The ubiquity of compound lipids in cellular biology places their metabolic regulation as a key factor in health and disease. Indeed, biological processes such as autophagy and vesicle secretion are intimately tied to lipid metabolism^11–13^, suggesting that measurements of compound lipid metabolic flux can be informative for cell and tissue health.

Given the bioenergetic costs of lipid synthesis as well as their intermediate molecular size relative to small molecule metabolites and proteins, compound lipids are synthesized *de novo* and extensively recycled by organisms^13–15^. At the same time, exogenous lipids are consumed, metabolized, and incorporated into cellular lipid pools such that membrane biochemistry can also reflect the cellular microenvironment or diet^16–20^. This inherent plasticity confers some resilience when cells have limited resources but also allows environmental lipids to influence membrane biology^21,22^. The vast, interconnected, network of enzymes responsible for headgroup and acyl-chain synthesis, elongation, compound lipid assembly, and catabolism pose a challenge to the identification of disease mechanisms associated with lipid homeostasis. However, recent advances in high-resolution mass spectrometry and computational metabolic flux analysis (MFA) are now enabling more reliable measurement of these processes.

^13^C MFA is the most advanced and reliable approach for quantifying biochemical fluxes in living organisms^23–27^. Culturing of cells with stable isotope-labeled nutrients generates complex labeling patterns or “isotopologue distributions” in lipids downstream of biosynthetic pathways. Beyond the generation and piece-wise assembly of compound lipids, these species are highly catabolized to facilitate the recycling and re-use of acyl-chains and headgroups in the lysosome, peroxisome, endoplasmic reticulum, Golgi, and mitochondria. Such metabolic exchange fluxes are best quantified using mechanistic models comprised of bi-directional and/or compartmentalized reactions within a comprehensive biochemical network^28^. Although commonly overlooked in favor of “rate limiting,” or unidirectional reactions within biochemistry texts, high exchange flux reactions promote the resilience of such systems to biochemical stressors and fluctuations in nutrient availability. For example, oxidative and reductive fluxes through isocitrate dehydrogenases support the use of different substrates for acetyl-coenzyme A provision in normoxia and hypoxia^29–31^. The malate-aspartate shuttle and pyruvate cycling similarly allow cells to negotiate anaplerosis and electron transfer within mitochondria, highlighting the importance of such metabolic exchange nodes. ^13^C MFA applications have typically addressed questions in central carbon metabolism and biopolymer synthesis^24,26,32^, with more targeted approaches used to examine lipid metabolism by examining distinct isotopologues^33^. As compound lipid metabolism becomes increasingly relevant in biology^34–36^, new tools are needed to extend MFA beyond these applications.

Here we develop and implement Compound Lipid MFA (CL-MFA) for the elucidation of reaction fluxes involved in lipid homeostasis, applicable to virtually any organism. This analytical and modeling framework leverages high-resolution mass spectrometry of intact compound lipids, metabolic networks encompassing compound lipid synthesis and recycling, and the INCA2.0 MFA^37^ platform to resolve fluxes across the lipidome. We demonstrate the isozyme specificity of a ceramide synthase inhibitor and the responsiveness of distinct compound lipids to serum availability. We further validate this approach in cultured cells and precision-cut lung slice cultures using non-small cell lung cancer (NSCLC) models that are liver kinase B1 (LKB1)-proficient or deficient. The LKB1 tumor suppressor controls lysosomal biogenesis and lipid recycling through AMP-activated protein kinase (AMPK) signaling^38,39^, and CL-MFA enables resolution of these downstream fluxes. This framework therefore enables reliable quantitation of metabolic fluxes across the lipidome in virtually any biological system, including homogenous cells in culture and precision-cut lung slice cultures cultured with stable isotope tracers, providing a versatile tool for researchers to assess the function of ∼1000 enzymes in lipid metabolism.

### Design, implementation, and assumptions

MFA modeling requires a stoichiometric network and isotopologue data to estimate relative fluxes and confidence intervals representing the metabolism of a biological system. For initial testing and implementation of CL-MFA, we developed a pipeline for generating metabolic networks that encompass both glycerolipid and sphingolipid metabolism **(Figure 1A).** To this end, we cultured the A549 NSCLC cell line in the absence of tracers and measured quantifiable lipid species of interest that were free of co-eluting, isobaric metabolites and displayed isotopologue distributions that aligned with calculated theoretical natural abundances **(Figure S1A)**. Using ultra-high pressure liquid chromatography coupled to high-resolution mass spectrometry, we identified a broad range of species across several lipid classes with confirmed MS2 fragments that were suitable for modeling, focusing on abundant species involved in membrane lipid homeostasis such as phosphatidylcholines, phosphatidylethanolamines, phosphatidylserines, and sphingolipids **(Table S1)**. For simplicity, we focused on *de novo* synthesized lipids with high enrichment and did not include species containing poly-unsaturated fatty acids (PUFAs), where peaks often contain numerous isobaric species. However, the network and data inputs can be readily tailored to incorporate specific lipids, pathways, tracers (e.g. isotopically labeled choline or PUFAs), and validated measurements depending on the question of interest.

**Figure 1.**
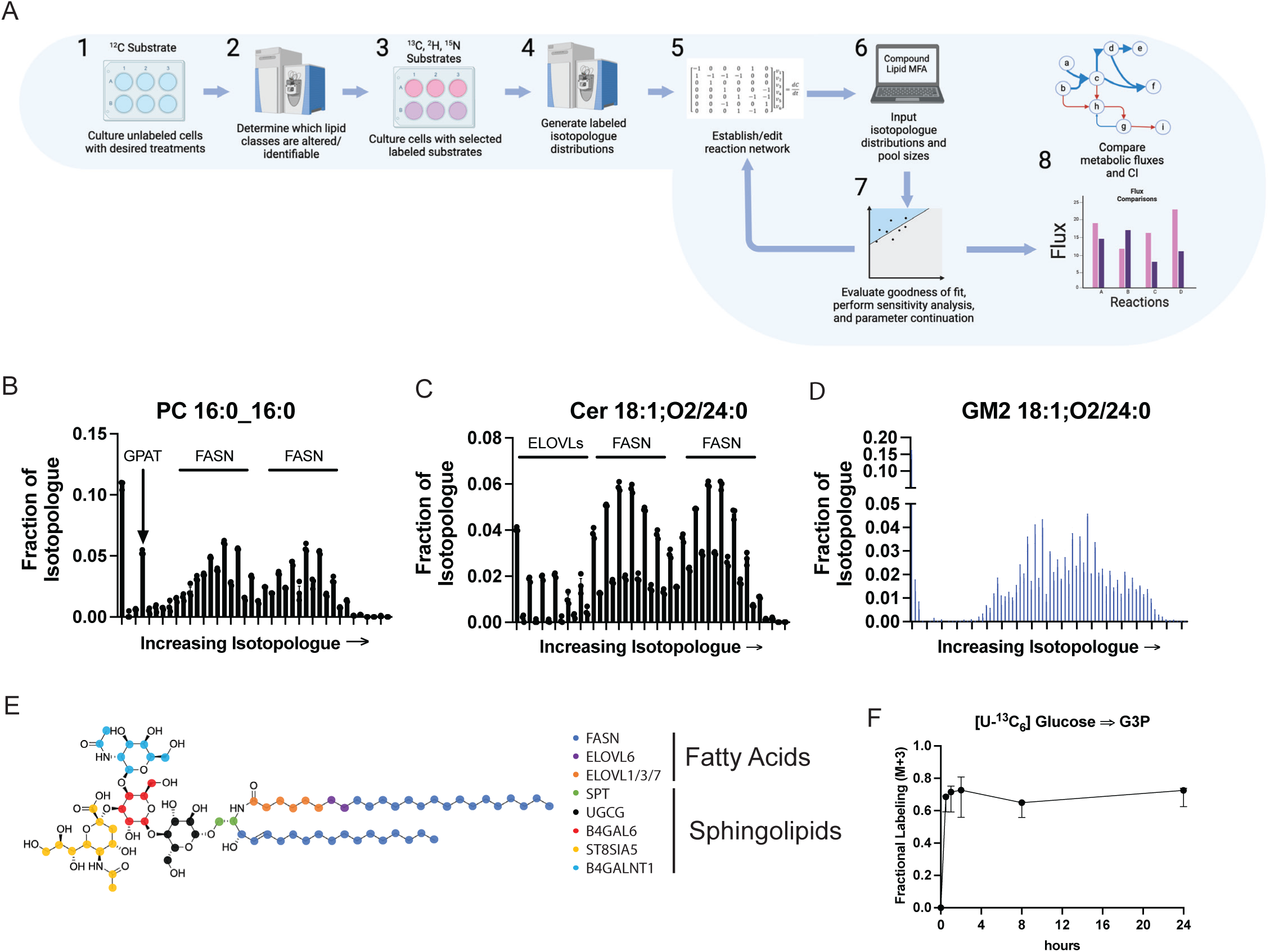
Design, implementation, and assumptions. (A) Workflow for the implementation of CL-MFA. (Created with Biorender.com) (B) Mass isotopologue distribution of PC 16:0_16:0 highlighting characteristic labeling patterns from FASN and GPAT flux from a [U-^13^C_6_]glucose tracing experiment (n=3). (C) Mass isotopologue distribution of Cer 18:1;O2/24:0 highlighting characteristic labeling patterns from FASN and ELOVL1/6 flux from [U-^13^C_6_]glucose tracing experiment (n=3). (D) Mass isotopologue distribution of GM2 18:1;O2/24:0 from a [U-^13^C_6_]glucose tracing experiment (n=3). (E) Depiction of GM2 18:1;O2/24:0 with colored carbon atoms matching enzymatic flux with potential labeling in the fractional labeling shown in Figure 1D. (F) Fractional labeling (M+3) of glycerol-3P from [U-^13^C_6_]glucose over 24 hours from A549 cells cultured in 10% FBS (n=3).

Next, we conducted tracer studies using [U-^13^C_6_]glucose, which yielded comprehensive labeling patterns after 24 hours for lipids with glycerol backbones and *de novo* synthesized or elongated fatty acids. Enrichment within compound lipids yields information on various enzymes involved in their biosynthesis and catabolism. Exemplary isotopologue distributions for phosphatidylcholine (PC), ceramide (Cer), and the ganglioside HexNAcNeuAcHex2Cer (GM2) are depicted in **Figures 1B-D** for reference. For GM2 18:1;O2/24:0, we highlight the atoms associated with enzymatic fluxes to headgroup biosynthesis, fatty acid synthase (FASN) activity, acylation of long-chain bases (LCBs) by ceramide synthase 2 (CERS2), elongation by fatty acid elongases (ELOVLs), and glycosylation using distinct colors (**Figures 1E)**. While the isotopologue patterns are visually distinct for some compound lipids like PCs **(Figure 1B)** and ceramides **(Figure 1C)**, as the number of building blocks increases, these data become increasingly more complex **(Figure 1E)**. By defining the atom transitions within a reaction network encompassing compound lipid synthesis and catabolism, the isotopologue distributions noted above can be used to estimate fluxes within the system. While we observed robust labeling in abundant compound lipids when culturing with [U-^13^C_6_] glucose, limited enrichment was observed in free sphingosine (SO) and sphinganine (SA) in whole cell lysates **(Figures S1B-C)**. These species are derived from catabolism of “cold” membrane sphingolipids within the lysosome and cytosol^40–42^. Thus, to observe robust flux through the serine palmitoyltransferase (SPT) complex, responsible for the *de* novo synthesis of LCBs from serine and palmitoyl-CoA, we cultured A549 cells in the presence of [U-^13^C_3_]serine. This tracer readily labeled dihydroceramide (DHCer), ceramide, and sphingomyelin pools in cell lysates while minimally labeling fatty acids and lipids aside from phosphatidylserine (PS) and phosphatidylethanolamine (PE) when serine biosynthesis was not observed **(Figures S1D-E)**. The inclusion of parallel serine tracing data facilitated measurement of *de novo* LCB synthesis versus LCB salvage from sphingomyelin, which is important for plasma membrane signaling and ceramide homeostasis^8^.

Next, we incorporated isotopologue data into a mechanistic ^13^C MFA model using INCA, which enabled estimation of fluxes throughout the network and calculation of confidence intervals for each via sensitivity analysis^43^. In the case of compound lipids, precursor pools (e.g. glycerol-3-phosphate, G3P, acetyl-CoA, serine) are assumed to be at pseudo-steady state showing relatively constant labeling **(Figures 1F, S1F)**, and the contribution of tracer to these pools is determined as the “D value.” Enrichment of compound lipids in biomass does not reach steady state and is calculated as fractional synthesis or *g(t)*^32,44^. These assumptions and solutions are routinely used for *in vivo* and *in vitro* flux measurements such as isotopomer spectral analysis (ISA) or mass isotopomer distribution analysis (MIDA)^44,45^. The reaction network and flux estimations for this study are presented in **Tables 1-14**. Separate compartments of acetyl-CoA for the cytosol and endoplasmic reticulum were defined to deconvolute FASN versus elongation flux taking advantage of the differential compartmentalization of FASN and the ELOVL enzymes. Overall, by incorporating isotopologue data from diverse lipid classes, we could resolve key fluxes in compound lipid biosynthesis, recycling, and interconversions between lipid classes.

For each CL-MFA analysis, we either 1) fix the final measured metabolite pool to 100 nanomoles without constraining intermediate or labeling pools or 2) use measured pool sizes to quantify nanomolar flux values necessary to maintain the observed membrane lipid pools. Finally, parameter continuation was used to determine the 99% confidence interval flux bounds for each reaction. Below, we apply CL-MFA to quantify fluxes across the lipidome within NSCLC models, exploiting distinct tumor genetics to highlight the ability of this tool for quantifying lipid synthesis and salvage fluxes within the ER, Golgi, and lysosomal network.

## Results

### Lipid availability strongly impacts acetyl-CoA and FASN flux in tissue culture

First, we determined if CL-MFA could resolve flux changes driven by culturing cells in media with reduced lipid availability. We performed parallel [U-^13^C_6_]glucose and [U-^13^C_3_]serine tracing experiments for 24 hours in the presence of either 1% or 10% FBS. We then incorporated key isotopologue distributions for separate reaction networks focused on sphingolipid and glycerolipid homeostasis **(Figures 2A, S2A, Tables 1-4)**. For analysis of sphingolipid metabolism, we built a reaction network that encompassed the *de novo* biosynthesis and recycling of SM 18:1;O2/24:0 and incorporated isotopologue data for SPB 18:1;O2, Cer 18:0;O2/24:0, Cer 18:1;O2/24:0, and SM 18:1;O2/24:0 generated from parallel tracer experiments using [U-^13^C_6_]glucose and [U-^13^C_3_]serine. Serum depletion increased the contribution of glucose to cytosolic acetyl-CoA and ER acetyl-CoA, and FASN flux increased in cells cultured with 1% FBS **(Figure 2A**, **Tables 1-2 R1, R9)**. However, this treatment caused a reduction in elongation flux from palmitate (FA 16:0) to lignoceric acid (FA 24:0) **(Figure 2B)**. In both conditions, fatty acid elongation through ELOVL1 and ELOVL6 was responsible for more than 80% of the lignoceric acid used to maintain sphingolipid pools (**Figure 2B**), indicating these species are highly synthesized. In contrast to acetyl-CoA and fatty acid biosynthesis, the contributions of *de novo* **(Tables 1-2, R27)** and salvage fluxes **(Tables 1-2 R28)** through ceramide synthase 2 (CERS2), the CERS isoform that generates either Cer 18:0;O2/24:0 and Cer 18:1;O2/24:0 depending on which LCB is the used substrate, were unchanged to maintain sphingolipid pools and there was no change to the turnover rate of the SM pool **(Figures S2B-C)**. While we observed an increase in the relative abundances of SA and DHCer at 1% FBS, the SO, Cer and SM pools were generally unchanged **(Figure 2C)**.

**Figure 2.**
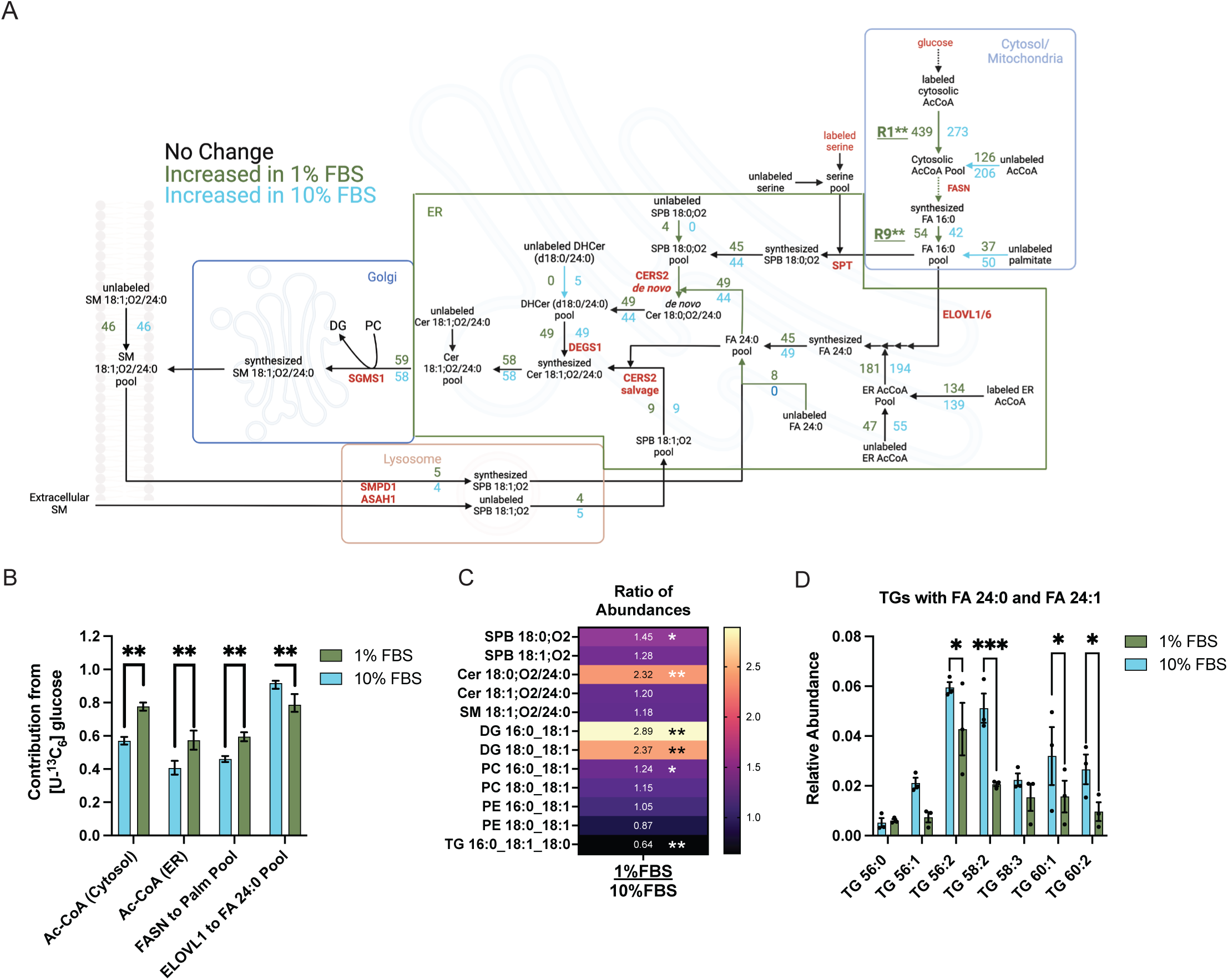
Lipid availability strongly impact acetyl-CoA and FASN flux in tissue culture. (A) Schematic of SM 18:1;O2/24:0 MFA with the net fluxes estimated from ^13^C CL-MFA for A549s cultured in 1% and 10% FBS for 24 hours. Fluxes represent the nanomoles needed to maintain a pool of 100 nanomoles of SM 18:1;O2/24:0. Black arrows represent non-overlapping confidence intervals while colored arrows represent higher flux values with non-overlapping 99% confidence intervals in green or blue for cells cultured in 1% FBS or 10% FBS, respectively. (Created with Biorender.com) (B) Percent contribution of [U-^13^C_6_]glucose to the different acetyl-CoA pools or enzymatic flux from isotopically labeled atoms to maintain the fatty acid pool. (C) Heatmap showing ratio (1% FBS: 10% FBS) of abundances between cells cultured in 1% FBS and 10%FBS for 24 hours (n=3). (D) Relative abundances of triacylglyceride molecules for which MS2 fragmentation confirms the presence of VLCFA, FA 24:0 and FA 24:1 (n=3). Relative abundance is calculated by normalizing to internal standard specific to lipid class. In Figure 2B, data shown are ratio of newly synthesized metabolite divided by the total active pool of metabolite as estimated by ^13^C MFA with 99% confidence intervals. In Figure 2E, data are mean ± standard error of mean (SEM) and were analyzed using an independent t-test (A,D). *p < 0.05, **p < 0.01, *** p< 0.001.

Next, we generated a glycerolipid reaction network to examine how FBS restriction impacted these reactions **(Figure S2A, Tables 3-4)**. We observed the same effect on cytosolic acetyl-CoA and FASN contributions to downstream metabolites **(Figures S2D)** as described above. While FASN flux was increased in cells cultured in 1% FBS, the fractional synthesis of glycerophospholipids was not changed, though cells cultured in 10% FBS had an increase in triacylglyceride (TG) turnover **(Figure S2E)**. To look more deeply at VLCFA metabolism, we quantified several TGs for which we could identify both lignoceric acid (FA 24:0) and nervonic acid (FA 24:1) in the MS2 data **(Figure 2D)**. Consistent with the predicted flux, we observed a reduction in FA24:0 and FA24:1 containing TGs in cells cultured in 1% FBS and an increase in both the diacylglycerides (DGs) DG 16:0_18:1 and DG 18:0_18:1 **(Figure 2C)**. On the other hand, PC and PE abundances were unchanged with FBS modulation despite their active turnover **(Figure 2C)**. Collectively, these data indicate that 1) acyl-chain synthesis and uptake are regulated distinctly from elongation and compound lipid turnover, and 2) CL-MFA can resolve specific changes in sphingolipid, phospholipid, and neutral lipid turnover within cells.

### LKB1 regulates long-chain base recycling through the lysosome

A549 cells lack the LKB1 tumor suppressor and exhibit aberrant AMPK signaling, which influences lipid homeostasis and lysosomal metabolism^32,46–48^. This lack of metabolic plasticity may account for the high rates of *de novo* biosynthesis observed in LKB1-deficient cancers^49,50^. To determine whether CL-MFA could resolve potentially dysregulated salvage fluxes in NSCLC, we stably expressed wild-type LKB1 (+LKB1) or an empty vector (+EV) in A549 cells **(Figure S3A)** and cultured each line in 10% FBS containing either [U-^13^C_6_]glucose or [U-^13^C_3_]serine for 24 hours. Cellular lipids were then extracted and measured for isotope enrichment, and these data were analyzed using the sphingomyelin **(Figure 3A**, **Tables 5-6)** and glycerolipid **(Tables 7-8)** networks described above.

**Figure 3.**
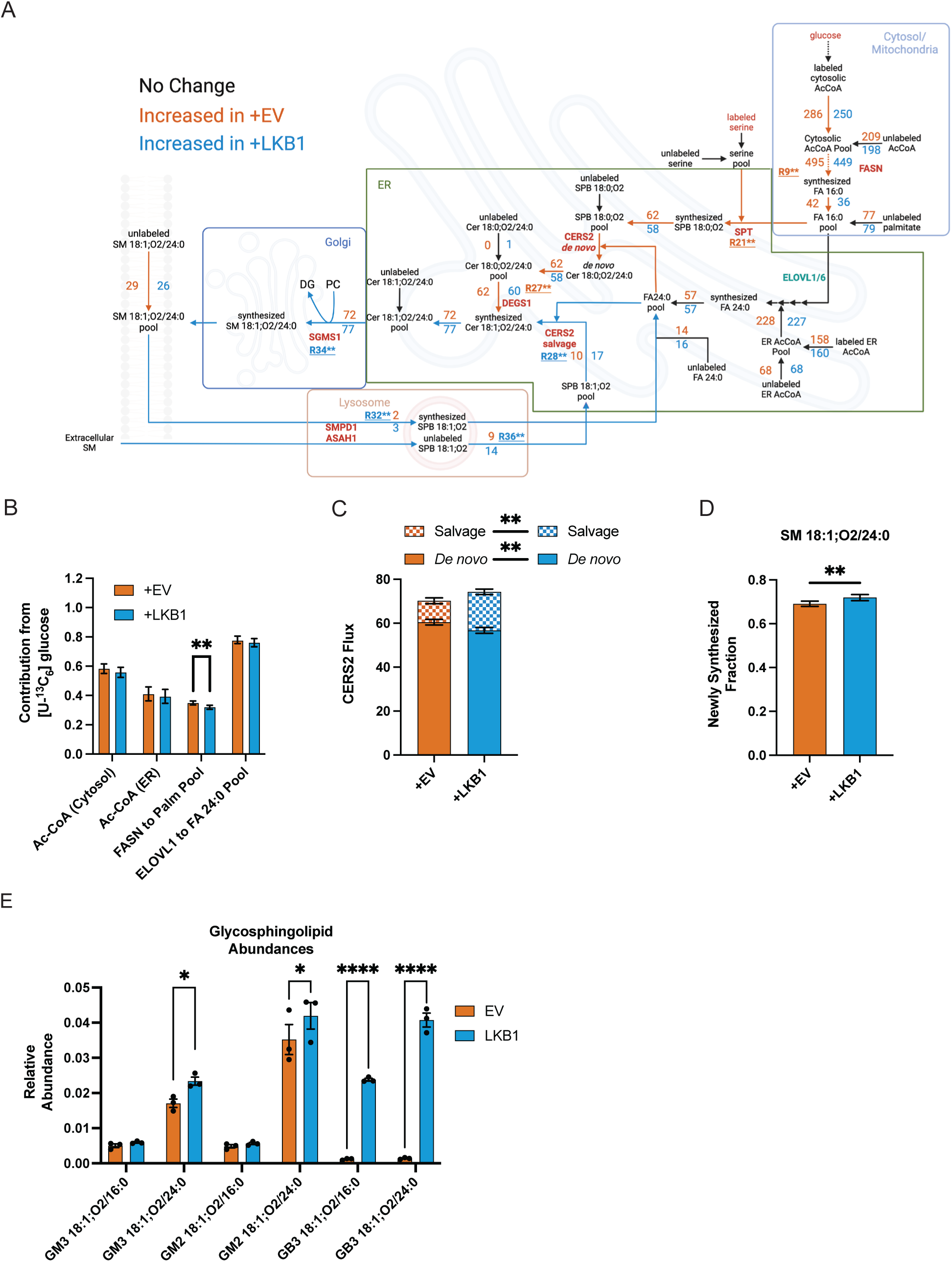
LKB1 regulates long-chain base recycling through the lysosome. (A) Schematic of SM 18:1;O2/24:0 MFA with the net fluxes estimated from ^13^C CL-MFA for A549 cells expressing an empty vector (+EV) or functional LKB1 (+LKB1) in 10% FBS for 24 hours. Fluxes represent the nanomoles needed to maintain a pool of 100 nanomoles of SM 18:1;O2/24:0. Black arrows represent non-overlapping confidence intervals while colored arrows represent higher flux values with non-overlapping 99% confidence intervals in orange or blue for cells expressing an empty vector or LKB1, respectively. (Created with Biorender.com) (B) Percent contribution of [U-^13^C_6_]glucose to the different acetyl-CoA pools or enzymatic flux from isotopically labeled atoms to maintain the fatty acid pool. (C) Comparison of CERS2 flux separated by the LCB used to generate the product of DHCer or Cer depending on sphinganine or sphingosine being the LCB used, respectively. (D) Newly synthesized fraction (g(t)) of SM 18:1;O2/24:0 after 24 hours of culture. (E) Relative abundances of glycosphingolipids in +EV and +LKB1 A549 cell lines (n=3). Relative abundance is calculated by normalizing to internal standard specific to lipid class. In Figures 3B, 3D, data shown are ratio of newly synthesized metabolite divided by the total active pool of metabolite as estimated by ^13^C MFA with 99% confidence intervals. In Figure 3C, data shown are estimated fluxes through CERS2 using sphinganine or sphingosine, simplified as *de novo* and salvage LCBs respectively, with 99% confidence intervals. In Figure 3E, data are mean ± standard error of mean (SEM) and were analyzed using an independent t-test (A,D).** p<0.01 for non-overlapping 99% confidence intervals, **** p< 0.0001.

We observed no difference in growth or the contribution of glucose to acetyl-CoA or glycerol-3 phosphate pools used for CL-MFA **(Figures 3B, S3C)**. However, the re-introduction of functional LKB1 in A549 cells decreased the contribution of FASN to the palmitate pool and the synthesis of glycerolipids containing palmitate, namely PC 16:0_18:1 and PE 16:0_18:1 **(Figures 3B, S3C-D)**. While the overall flux through CERS2 was comparable between these cells, the contribution of LCB salvage to sphingolipid pools was increased in +LKB1 cells **(Figures 3A, 3C, Table 5-6 R28)**, while *de novo* LCB synthesis to SL pools was increased in +EV cells **(Figures 3A, 3C, Table 5-6 R27)**. Specifically, free SO recycled from membrane sphingolipids and media sphingolipids contributed more nanomoles and a greater percentage to ceramide synthesis in +LKB1 cells versus +EV cells **(Figure 3A**, **Tables 5-6 R32, R36).** The increase in SM breakdown and salvage of SO resulted in increased SM turnover in the plasma membrane **(Figure 3D**, **Tables 5-6 R34)**. We also observed a reduction in Cer 18:0;O2/24:0 and SM 18:1;O2/24:0 and an increase in PC 16:0_18:1 and PC 18:0_18:1 pools, suggesting a shift towards more palmitate and stearate (FA 18:0) salvage and less *de novo* synthesis of sphingolipids in +LKB1 cells as compared to +EV cells **(Figure S3E)**. These data suggest that LKB1-proficient cells exhibit higher plasma membrane lipid exchange, consistent with its role in AMPK signaling and lysosomal biology.

We also measured glycosphingolipid (GSL) abundances in each cell line and observed marked increases in the enrichment and abundance of globosides (GB) in +LKB1 cells **(Figure 3E)**, highlighting links between AMPK signaling and GSL metabolism. These membrane lipids influence receptor-mediated signaling, cell adhesion, and cell-cell interactions^2,11,51–53^, so their metabolism is intimately tied to cell identity and phenotype.

To further evaluate the ability of CL-MFA to resolve synthesis and salvage fluxes across cell types, we next compared sphingolipid fluxes in parental A549 cells and the H1299 cell line, which expresses functional LKB1 alleles^54^ **(Figures S3F-H, Table 9)**. H1299 cells exhibited reduced palmitate and lignoceric acid (FA 24:0) synthesis as well as higher lipid salvage fluxes compared to A549 cells. In contrast, A549 cells showed higher flux through biosynthetic pathways spanning FASN through SPT **(Tables 1 and 9, R9, R12, R21, R28, R32, R36)**. These observations provide additional evidence that CL-MFA can resolve changes in lipid homeostasis in cultured cells **(Figures S3E-G)**.

### Application of CL-MFA to precision-cut lung slice culture

Next, we examined whether CL-MFA could resolve the same flux changes in a more complex, physiologically-relevant microenvironment, applying precision-cut lung slice (PCLS) culture to mice bearing NSCLC tumors^55,56^. We initiated lung tumors in Kras^LSL-G12D/+^; LKB1^flox/flox^ (KL) and Kras^LSL-G12D/+^; Trp53^flox/flox^ (KP) via intratracheal intubation of lentiviral Cre^49,57^. Approximately 16 weeks after tumor initiation, lungs were inflated with agarose and prepared for slicing and sub-culture as described in the methods. After overnight recovery in fresh medium, slices were cultured in media containing either [U-^13^C_6_]glucose or [U-^13^C_3_]serine for 24 hours **(Figure 4A)**. Finally, slices were processed and analyzed for CL-MFA as described above.

**Figure 4.**
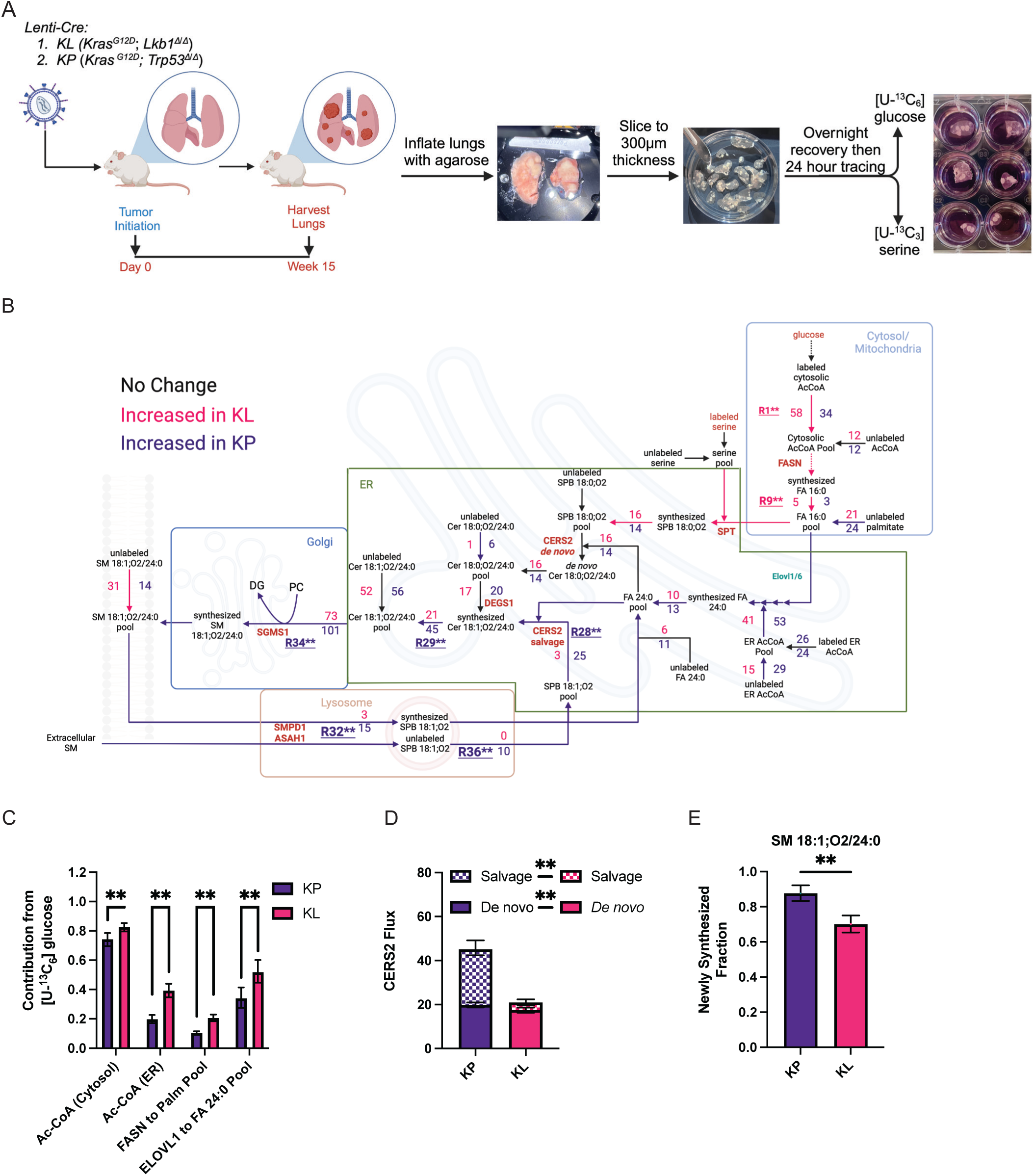
Application of CL-MFA to precision-cut lung slice culture. (A) Schematic of the generation of precision-cut lung slices from GEMMs harboring KL and KP mutations. (Created with Biorender.com) (B) Schematic of SM 18:1;O2/24:0 MFA with the net fluxes estimated from ^13^C CL-MFA for precision-cute lung slices harboring KL and KP tumors. Fluxes represent the nanomoles needed to maintain a pool of 100 nanomoles of SM 18:1;O2/24:0. Black arrows represent non-overlapping confidence intervals while colored arrows represent higher flux values with non-overlapping 99% confidence intervals in magenta or purple for tumors harboring KL and KP mutations, respectively. (Created with Biorender.com) (C) Percent contribution of [U-^13^C_6_]glucose to the different acetyl-CoA pools or enzymatic flux from isotopically labeled atoms to maintain the fatty acid pool. (D) Comparison of CERS2 flux separated by the LCB used to generate the product of DHCer or Cer depending on sphinganine or sphingosine being the LCB used, respectively. (E) Newly synthesized fraction (g(t)) of SM 18:1;O2/24:0 after 24 hours of culture. Relative abundance is calculated by normalizing to internal standard specific to lipid class. In Figures 4C, 4E, data shown are ratio of newly synthesized metabolite divided by the total active pool of metabolite as estimated by ^13^C MFA with 99% confidence intervals. In Figure 4D, data shown are estimated fluxes through CERS2 using sphinganine or sphingosine, simplified as *de novo* and salvage LCBs respectively, with 99% confidence intervals.** p<0.01 for non-overlapping 99% confidence intervals

Model output results largely reflected the expected changes in flux across these tumor genotypes **(Figure 4B**, **Table 10-11)**. Lung slice cultures from mice bearing KP tumors exhibited higher salvage flux and reduced contribution of glucose to acetyl-CoA pools fueling FASN **(Figure 4B-C, Tables 10-11 R1, R9, R12)**. In contrast, KL tumor slice cultures revealed higher biosynthetic flux through FASN and SPT, lower recycling of LCBs and an overall decrease in SM turnover **(Figures 4B-D**, **Tables 10-11 R32, R34, R36)**. These data suggest that membrane lipid turnover is significantly higher in KP tumors compared to KL tumors, which are unable to effectively regulate AMPK signaling and lysosomal biogenesis^39,49^. Notably, these fluxes are not resolvable through systemic approaches such as administration of ^2^H_2_O, as we have previously tested in similar models^50^. Finally, we observed that KL tumors maintained higher pools of sphingolipids in the SM 18:1;O2/24:0 synthesis axis, with significant increases in SA, DHCer and SM pools **(Figures S4A-D)**. These abundance data correlate with the elevated SPT flux and reduced salvage fluxes resolved by CL-MFA in KL tumor-bearing PCLS cultures **(Figure 4B, Tables 10-11 R9, R32, R36)**. Our approach can therefore elucidate flux changes in microenvironments containing diverse cell types that are likely to support lipid metabolism through distinct enzymatic activities, systems that better recapitulates tissue- or tumor-level homeostatic processes that are more physiological relevant than cultured cell lines.

### Mammalian tissue culture promotes reliance on FASN flux

In comparing our flux measurements in PCLS tumors **(Figure 4A)** to those generated in cell lines **(Figures S3F-H, Tables 1 and 9),** we noted that FASN and ELOVL1 fluxes were significantly lower in slice culture versus A549 and H1299 cell culture. Furthermore, slice cultures were maintained in 2% FBS versus the commonly used 10% FBS employed for tissue culture. We therefore hypothesized that even 10% FBS may be highly non-physiological compared to *in vivo* microenvironments, driving elevated lipid synthesis in proliferating cell cultures. To directly address this question, we applied CL-MFA using parallel [U-^13^C_6_]glucose and [U-^13^C_3_]serine tracing for 24 hours in A549 cells in the presence of 10% or 20% FBS, which showed relatively similar growth rates over this period of culture time **(Figure S5A)**. Notably, we observed a marked decrease in the contribution of glucose to acetyl-CoA as well as flux through FASN in the presence of 20% FBS **(Figure 5A)**. This reduction in *de novo* palmitate synthesis was accompanied by an increase in unlabeled palmitate used for elongation and compound lipid synthesis, as highlighted by the isotopologue distribution of Cer 18:1;O2/24:0 **(Figure 5B)**. In contrast to this marked decrease in FASN flux, ELOVL1 activity was only slightly reduced, highlighting the importance of VLCFA availability and synthesis toward cell growth regardless of FBS availability^58^. We observed no changes in lipid abundances within the SM reaction network, highlighting the increased uptake of SM to balance the reduction in synthesis, and the abundances of PC 16:0_18:1 and PC 18:0_18:1 were increased **(Figure 5C).** Furthermore, supplementation of 20% FBS had minimal impacts on growth, CERS2 flux, or the overall turnover of sphingomyelin **(Figures S5A-C)**, and the contribution of serine towards LCB synthesis was only mildly impacted **(Figure S5D).** Taken together, these findings highlight a “separation” between cytosolic fatty acid metabolism and the biosynthesis of complex, bioactive lipids that require additional modifications such as elongation, desaturation, or headgroup alterations occurring in the ER and Golgi. Our results also suggest that lipid availability and fatty acid metabolism are poorly modeled in typical cell culture conditions that employ 10% FBS, which selects for cells with high ACC and FASN flux, highlighting the importance of the tumor microenvironment (TME) in supporting lipid homeostasis.

**Figure 5.**
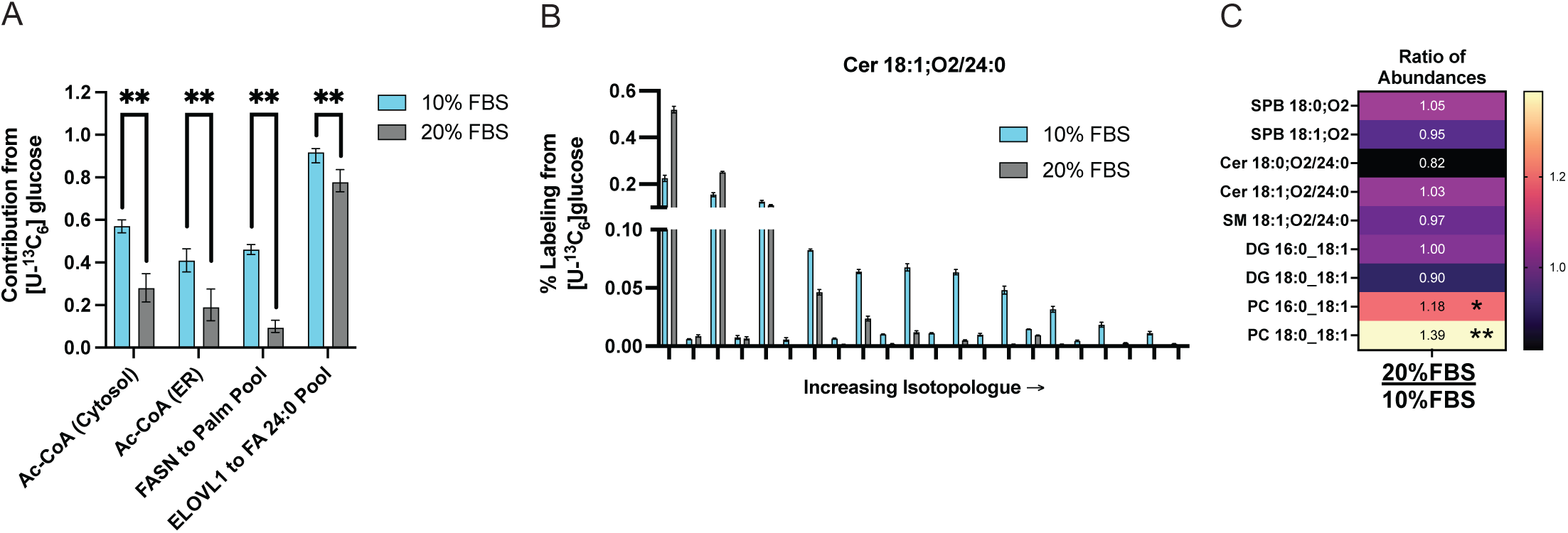
Mammalian tissue culture is inherently deficient in lipids. (A) Percent contribution of [U-^13^C_6_]glucose to the different acetyl-CoA pools or enzymatic flux from isotopically labeled atoms to maintain the fatty acid pool. (B) Mass isotopologue distributions of Cer 18:1;O2/24:0 from cells cultured in 10% and 20% FBS (n=3). (C) Heatmap showing ratio (20% FBS:10% FBS) of abundances between cells cultured in 20% FBS and 10%FBS for 24 hours (n=3). Relative abundance is calculated by normalizing to internal standard specific to lipid class. In Figure 5C, data are mean ± standard error of mean (SEM) and were analyzed using an independent t-test (A,D). * p< 0.05, ** p<0.01

### CL-MFA resolves isozyme specificity of drugs

Enzymes involved in lipid homeostasis are highly druggable clinical targets^58–64^, but promiscuity and redundancies in these pathways complicate their analysis. Pharmacological agents may also have varied affinity for isozymes, leading to off-target effects that could impact lipid metabolism more broadly. To assess the ability of CL-MFA to identify such mechanistic changes, we built a model encompassing both glycerolipid and sphingolipid reaction networks **(Figure 6A**, **Tables 13-14)** and examined flux changes induced by fumonisin B1 (FuB1), a commonly used “pan-inhibitor” of CERS isozymes. This broader analysis incorporated isotopologue data for sphingomyelin, glycosphingolipid, phosphatidylcholine, phosphatidylethanolamine, and phosphatidylserine species into an integrated model to estimate fluxes across the network and quantify changes in flux through two CERS isoforms, CERS6 and CERS2, responsible for the acylation of SA and SO with either palmitate or a VLCFA, respectively.

**Figure 6.**
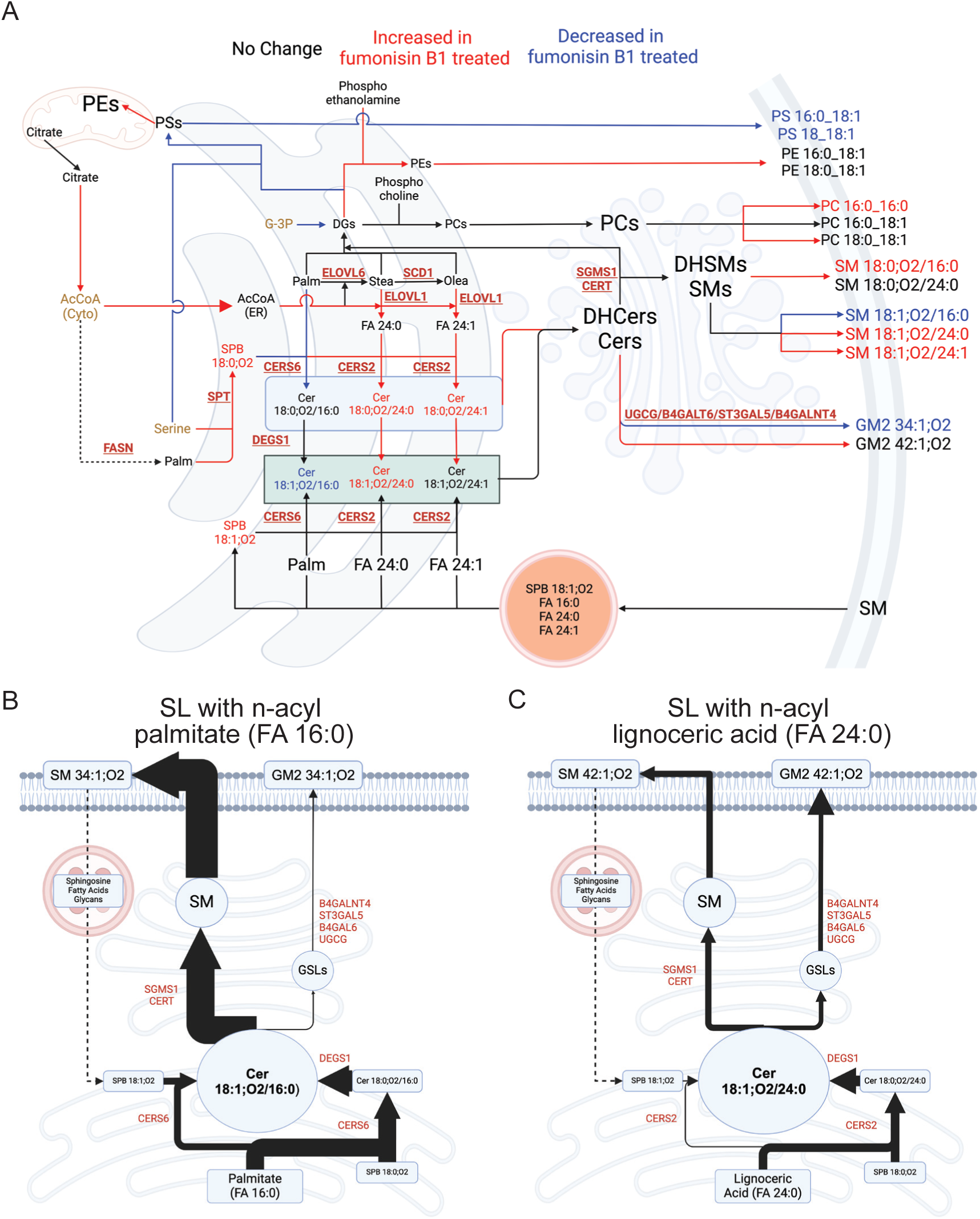
Application of CL-MFA to precision-cut lung slice culture. (A) Schematic of CL-MFA with the net fluxes estimated from ^13^C CL-MFA for cells cultured in 20%FBS with vehicle or 2uM fumonisin B1. Fluxes represent the nanomoles needed to maintain the measured pool sizes for the final compound lipids detailed on the plasma membrane. Black arrows represent non-overlapping confidence intervals while colored arrows represent higher flux values with non-overlapping 99% confidence intervals in blue or red for A549s cells cultured with vehicle or fumonisin B1, respectively. (Created with Biorender.com) (B) Depiction of Cer 18:1;O2/16:0 as the hub for sphingomyelin synthesis or glycosphingolipid synthesis depending on trafficking in A549s cells cultured with vehicle in 20%FBS. (C) Depiction of Cer 18:1;O2/24:0 as the hub for sphingomyelin synthesis or glycosphingolipid synthesis depending on trafficking in A549s cells cultured with vehicle in 20%FBS. Thickness of arrows for Figures 6B-C represent the flux for each enzyme.

A549 cells were cultured in the presence of 2μM FuB1, a concentration that does not inhibit cell growth (**Figure S6A**). Strikingly, we observed divergent effects on reactions catalyzed by CERS6 versus CERS2. As expected, FuB1 potently reduced flux through CERS6 observed by a reduction in Cer 18:0;O2/16:0, Cer 18:1;O2/16:0, and SM 18:1;O2/16:0 synthesis, reduced pools of these sphingolipids and increased SPB 18:0;O2 and SPB 18:1;O2 abundances **(Figures 6A, S6B-C, Tables 13-14 R88, R92, R95, R96, R98, R147, R149)**. However, we observed a marked increase in flux through CERS2 along with elevated levels of sphingolipids containing the VLCFAs lignoceric acid and nervonic acid **(Figures 6A, S6B-C, Tables 13-14 R22, R39, R44, R47, R108, R111, R160, R162)**. Similar changes were observed in pool sizes and synthesis fluxes to GM2 gangliosides **(Figures 6A, S6B-C, Tables 13-14 R160, R162)**. To maintain the increased flux through CERS2 and reliance on VLCFAs, cells treated with FuB1 increased VLCFAs biosynthetic flux catalyzed by ELOVL1 **(Figure 6A**, **Tables 13-14, R1, R5, R22, R100)**. These changes in CERS isozyme fluxes caused further alterations throughout the network. For example, with reduced incorporation of palmitate into Cer 18:1;O2/16:0 and SM 18:1;O2/16:0, we detected a compensatory increase in PC 16:0_16:0, PC 16:0:18:1, and PE 16:0_18:1 and PE 18:0_18:1 synthesis, suggesting reduced palmitate containing sphingolipid biosynthesis drives increases of select phospholipids species**(Figures S6B-C, Tables 13-14 R52, R54, R68, R70)**. On the other hand, there was a reduction in the PS 16:0_18:1 and PS 18:0_18:1 pools along with their biosynthesis but an increase to PS to PE conversion which occurs in the mitochondria **(Figure 6A**, **Table 13-14 R66, R82)**.

Lastly, we used the model output to compare how short-chain (Cer 18:1;O2/16:0) or very long-chain (Cer 18:1;O2/24:0) ceramides were partitioned in downstream reactions to sphingomyelin and glycosphingolipids in A549 cells cultured in 20% FBS. While these components of lipid ordered domains are synthesized from the same ceramide pool, their fate is dependent on traffic to the Golgi by ceramide transfer protein (CERT) or coat protein complex II (COPII)-mediated vesicular flux^65–67^. We observed that ceramides containing palmitate as the n-acyl chain were converted to sphingomyelin at much higher rates than glycosphingolipids **(Figure 6B, Tables 13-14 R94, R137)**. On the other hand, flux of ceramides containing lignoceric acid (FA24:0) were evenly partitioned to sphingomyelin and GM2 gangliosides **(Figure 6C**, **Table 13-14 R44, R150)** These results collectively highlight the ability of CL-MFA to quantify fluxes through critical nodes of lipid homeostasis and capture the topology of lipid metabolism beyond the FASN-ACC axis and pool size changes.

## Discussion

Here we demonstrate the ability of CL-MFA to untangle the complexity of lipid metabolic pathways, providing a quantitative, systems-based approach for studying lipid homeostasis. For decades, stable isotope tracing and downstream modeling have provided key insights into biopolymer synthesis in cells, animals, and patients, highlighting critical tissue sites for lipogenesis in numerous disease contexts^25,44,68–70^. The labeling information and network analysis outlined here can provide quantitative information on headgroup and acyl-chain flux across numerous classes of compounds lipids. In turn, one can identify the most active (or inactive) lipid-metabolizing enzymes within a given biological system, including recycling pathways that may span several organelles or cell types as well as core pathways such as the Lands cycle. While custom tracers or wash-out approaches can facilitate measurement of fluxes associated with compound lipid uptake, catabolism, and recycling, their applications are costly and technically challenging^71^. More accurate and sensitive high-resolution mass spectrometry technology facilitate this method, providing unprecedented detail on lipid homeostasis^52,72,73^. This framework enables quantification of lipid metabolism beyond the established ACLY-ACC-FASN axis of reactions which have been the focus of most lipid flux studies to date. Our comparisons of lipid metabolism in PCLS versus cell culture in media with different serum supplementation also highlights the non-physiological nature of tissue culture. Lipids are readily available from the microenvironment (e.g. tumor interstitial fluid)^74^ or circulation *in vivo*, and our results suggest that lipid synthesis becomes highly active in less than 24 hours of culture in 10% FBS. These findings contrast observations of “polar” substrate concentrations in typical culture media that are present at supraphysiological levels^75,76^, such that tissue culture is inherently deficient in lipids relative to *in vivo* conditions. This result may have profound implications on the metabolism of cells adapted to *in vitro* culture and data generated under these conditions^77^.

The tumor microenvironment contains diverse cell types, and this heterogeneity facilitates metabolic crosstalk and exchange of lipids in addition to that provided by circulating lipoproteins^35,78–82^. This complex environment provides tumors with an array of pathways to support proliferation. Using PCLS culture, our approach successfully quantified changes in lipid salvage and biosynthesis in NSCLC tumors with specific genotypes. Indeed, autophagy, lipophagy, and other salvage pathways are emerging as critical mediators of tumor progression, neurodegeneration, and aging^13,14,83^, so precise quantitation of these pathways is becoming increasingly important in biology. The redundancy and flexibility of such exchange fluxes and metabolite interconversions provides an innate biochemical resilience that is necessary for cell survival. Now, we can quantitatively measure these phenomena within lipid metabolism.

While *in vitro* culture conditions will never perfectly recreate the tumor microenvironment, the molecular detail and tunability afforded by tissue culture CL-MFA will be an invaluable tool for mechanistic studies that identify gene function. Many genes encoding lipid metabolizing enzymes remain unannotated, and it remains challenging to resolve the distinct functionalities of tissue-specific isozymes or disease variants for key enzymes in lipid metabolism. This issue is particularly relevant in characterizing the function of sphingolipid enzymes linked to diseases such as ALS and sensory neuropathy^85–87^.

Finally, the application of CL-MFA will enable the testing of hypotheses focused on specific mechanistic questions while capturing broader changes in the lipidome at the systems-level. With higher resolution equipment becoming more available, the application of distinct stable isotope elements can add an additional layer of information. For example, application of [^13^C_6_]glucose and deuterated PUFAs, for examples, enables the modeling of *de novo* lipogenesis and uptake of essential fatty acids and their fates^88,89^. The use of spatial mass spectrometry imaging technology can, in turn, provide information on local flux changes (e.g. ACC-FASN) within a tissue^52,72^, and integration of these pipelines into CL-MFA may provide deeper information on lipid trafficking in distinct cell types. The comprehensive flux quantitation afforded by CL-MFA will prove invaluable for addressing such questions in the future, revealing the dynamic nature of membrane lipid metabolism.

## Limitations of the study

In the studies and models outlined here, we focused on the most abundant lipids that were actively synthesized and salvaged. The cellular lipidome is much more diverse and a full-scale cell lipidome MFA would require more expansive reaction networks. Therefore CL-MFA should be viewed as modular in nature such that focused studies on specific lipid pathways or sub-networks can be executed easily. We also predominantly conducted “steady-state” models assuming precursor enrichment is constant and estimating time-dependent fractional synthesis of lipid pools rather than performing kinetic, non-stationary MFA (INST-MFA). In test cases we found that sufficient information was available to resolve fluxes using steady-state modeling, and the computational time required for INST-CL-MFA modeling and parameter continuation is prohibitive for routine use. However, application of INST-CL-MFA is feasible if required to increase flux resolution further. Nevertheless, our results highlight the broad impacts of CL-MFA in assessing lipid metabolism to answer distinct biochemical questions spanning the lipidome.

## Supporting information

MFA results: A549 SL 1% FBS

MFA results: A549 SL 10% FBS

MFA results: A549 PL 1% FBS

MFA results: A549 PL 10% FBS

MFA results: A549 SL +EV 10% FBS

MFA results: A549 SL +LKB1 10% FBS

MFA results: A549 PL +EV 10% FBS

MFA results: A549 PL +LKB1 10% FBS

MFA results: H1299 SL 10% FBS

MFA results: KL lung Slice SL 2% FBS

MFA results: KP lung Slice SL 2% FBS

MFA results: A549 SL 20% FBS

MFA results: A549 20% FBS +veh

MFA results: A549 20% FBS + FuB1

Transition Tables for LCMS

## Acknowledgement

We thank all members of the Metallo Lab for helpful discussions. We acknowledge support from NIH grant R01CA234245 (to C.M.M.), the Lowy Medical Research Institute (to C.M.M.), the Mark Foundation for Cancer Research (to C.M.M. and R.J.S.), NIH grant R35CA35220538 (to R.J.S), the Salk NCI Cancer Center CCSG P30 CA013195, and an AHA-Allen Initiative in Brain Health and Cognitive Impairment award made jointly through the American Heart Association and The Paul G. Allen Frontiers Group: 19PABH134610000.

## Author contributions

C.M.M. and K.W.R. conceived and designed the study. K.W.R., E.L.A., and Y.H. performed in vitro mass spectrometry experiments and analysis. GEMMs were generated by H.M.G. and C.W.M. M.L.R. performed harvesting of lungs and performed the precision lung slicing. K.W.R. and M.L.R. cultured lung slices and separated tumors from lung adjacent tissue. H.M.G. and R.J.S. provided the A549 cell lines expressing an EV and LKB1. H.M.G. performed western blots. K.W.R., G.M. optimized LC-MS/MS parameters for lipid measurements. K.W.R. performed all reaction network design and CL-MFA in discussion with S.K. and C.M.M. K.W.R. analyzed all remaining data. C.M.M. and K.W.R. guided experimental design and analysis. K.W.R and C.M.M wrote the manuscript with input from all authors.

## Declaration of Interests

The authors declare no competing financial interests.

## Methods

### Cell culture experiments

A549 and H1299 cells were cultured in Dulbecco’s modified Eagle’s medium (DMEM, Gibco) with 1%, 10%, or 20% FBS and 1% penicillin/streptomycin (P/S). A549 cells were transduced with pBabe (empty vector; Addgene #1764) or pBabe-FLAG-LKB1 (Addgene #8592) LKB1 as previously described^90^. Cells with the integrated vectors were selected for using 400 ug/ml of Hygromycin in cell culture. Cell lines were confirmed to be free of Mycoplasma (Lonza LT07-318). All media were adjusted to pH = 7.3.

### Mouse studies

All animal studies were approved by the Salk Institute Institutional Animal Care and Use Committee. Mouse strains were maintained on FVB/N background. *KL* (*Kras^LSL-G12D/+^*;*Lkb1^flox/flox^*;*R26^LSL-Luc/LSL-Luc^*) and *KP* (*Kras^LSL-G12D/+^Trp53^flox/flox^*; *R26^LSL-Luc/LSL-Luc^*) mice have been described previously^49^. Lung tumors were initiated by the delivery of lentiviral Cre recombinase (4 × 10^5^ plaque-forming units per mouse) via intratracheal intubation as described previously^57^ Lung tissues were collected immediately for tissue slice culture upon observation of respiratory distress (15 – 18 weeks after tumor initiation).

### Precision cut lung slice culture

Low-melt agarose (LMA) was prepared prior to procuring lungs. 3% w/V low-melt agarose Type VII-A (Sigma, A0701) was added to a 0.9% NaCl solution, microwaved to dissolve, and placed into a 42°C water bath. A 3ml syringe tipped with a blunt 21-gauge needle was filled with 2ml of LMA and kept warm.

Mice were sacrificed and trachea was nicked with a 25-gauge needle. Warm LMA was pushed into the trachea using the pre-filled syringe. The lungs were slowly inflated with approximately 2ml of LMA, and the thorax packed with ice to solidify the agarose prior to withdrawing the needle from the trachea. The lungs were removed from the thorax en bloc, the lobes separated out and placed in chilled, sparged (with 95% O_2_/5% CO_2_ carbogen gas) slicing buffer. The lobes were superglued to a stage with the hilium facing downward, and 300μm sections cut on a Leica VT1200S with Vibracheck. Prior to incubation at 37°C, sections were kept continuously in chilled, sparged slicing buffer (Krebs-Henseleit buffer (Millipore Sigma, Cat# K3753). Lung slices were matched for size and tumor burden, placed into individual wells in a 12 well suspension culture plate (greiner bio-one Cellstar, #665 102) containing warmed, oxygenated resting media (2 ml, RPMI 1640 with 2%FBS (GIBCO^TM^ ThermoFisher Scientific, Cat# 11875093) and incubated overnight while on an orbital shaker.

Following an overnight recovery in RPMI, slices were washed three times with 0.9% saline and tracing DMEM media containing [U-^13^C_6_]glucose or [U-^13^C_3_]serine was added. After 24 hours, tracing media was collected and the lung slices rinsed 3 times with 0.9% (w/V) saline. After the final saline rinse was removed, the slices were quick frozen by placing the 12 well culture plate on top of a metal block immersed in a slurry of isopropanol and dry ice. Biopsy punches were used to isolate frozen disks of tumor and adjacent lung tissue for separate analysis.

### Cell proliferation and ^13^C tracing

Proliferation studies were performed on 6-well plates with an initial cell number of 50,000 per well for A549s parental and derivative cell lines. Cells were plated in growth media (10%FBS) and allowed to adhere for 24 hours before changing to the specified growth media. Cell counts were performed using the Incucyte SX5 for 72 hours after media change. Best fit exponential growth curves were calculated to fit the growth data and 95% confidence intervals for growth rates (hr^-1^) were calculated.

^13^C isotope tracing media was formulated using a custom Hyclone glucose-, amino acid-, and sodium pyruvate-free DMEM (Cytiva Life Sciences) supplemented with either 25 mM [U-^13^C_6_]glucose or 0.4 mM [U-^13^C_3_]serine while replacing the remainder amino acids or unlabeled glucose to achieve the necessary media composition and supplemented with the amount of FBS necessary to achieve the condition described in each experiment. Cultured cells were washed with 1 mL of phosphate-buffered saline (PBS) before applying tracing media for 24 hours as indicated in figure legends. Cells were cultured with vehicle or 2μM fumonisin B1 overnight prior to the application of tracing media.

### Reagents

[U-^13^C_6_]glucose (CLM-1396) and [U-^13^C_3_]serine (CLM-1574) were acquired from Cambridge Isotope Laboratories. Fumonisin B1 (BML-SL220) was purchased from Enzo Life Sciences and resuspended in DMSO.

### Antibodies

Antibodies from Cell Signaling Technologies (Danvers, MA USA) were diluted 1:1000 in sterile-filtered 1% BSA (Cat) in TBS-T 0.1%. Antibody targets included: LKB1 (#3047), p-ACC^S79^ (#3661), ACC (#3662), p-AMPKα^T172^ (#2535). Anti-actin (#A5441, Sigma Aldrich) was diluted 1:10000.

### Western Blots

Protein lysates were prepared in the following lysis buffer supplemented with protease (Roche, 11836153001) and phosphatase (Roche, 4906837001) inhibitors: 100 mM Tris-HCl (pH 7.4), 300 mM NaCl, 2% Triton X-100, 2 mM EDTA, 0.2% SDS, and 1% Sodium Deoxycholate. Cell lysates in prepared lysis buffer were homogenized by water bath sonication. Snap frozen micro-dissected lung tumors were homogenized on ice in 50 µL lysis buffer by mechanical dissociation followed by microtip sonication. Total protein input was normalized across samples by evaluating protein concentration by BCA (Thermo Fisher Scientific 23225). Denatured protein lysates were resolved on 4-12% Bis-Tris gels (Invitrogen, NP0335BOX) and transferred to PVDF membranes (Thermo Fisher Scientific, 88518). The resulting membranes were blocked in 5% milk in TBS-T 0.1%, incubated overnight at 4°C in diluted primary antibody, washed with TBS-T, incubated for one hour in secondary antibody diluted in in TBS-T plus milk, washed in TBS-T, and developed using Clarity Western ECL Subtrate (Bio Rad, 170-5060). Secondary antibodies were anti-rabbit (Millipore, AP132P) and anti-mouse (Millipore, AP124P).

### Metabolite extraction and GC-MS analysis

At the conclusion of the tracer experiment, media was aspirated. Then, cells were rinsed twice with 0.9% saline solution and lysed with 500 μL of ice-cold methanol. 10 nanomoles of norvaline was added to each sample along with compound lipid standards. Cells were scraped from the plate and lysates transferred to Eppendorf tubes. Samples were split in two. One portion was used for polar compound lipid analysis on LCMS and the other further extracted and separated to generate an aqueous layer containing polar metabolites for GCMS analysis and a hydrophobic phase used for analysis of neutral lipids on LCMS. Separation of polar and organic phases was performed by addition of 200uL of water and 400 μL of chloroform to the 200 μL of methanol left in the Eppendorf tube. Samples were then vortex and centrifuged at 21,000gs for 5 min at 4°C. The organic phase was collected and 2 μL of formic acid was added to the remaining polar phase which was re-extracted with 400 μL of chloroform. Combined organic phases were dried under nitrogen and used for neutral lipid analysis.

Polar lipid samples were centrifugation at 21,000 *g* for 15 min at 4°C. 300 μL of methanol were collected and evaporated under nitrogen and stored at −80°C until resuspended for analysis. 250 μL of the upper aqueous phase were transferred to a GC vial and dried at 4°C overnight. 250uL of lower organic layer was collected and evaporated under nitrogen at room temperature. Dried polar metabolites were processed for gas chromatography–mass spectrometry (GC-MS) as described previously by Cordes and Metallo^91^. Briefly, polar metabolites were derivatized using a Gerstel MultiPurpose Sampler (MPS 2XL). Methoxime–*tert*-butyldimethylchlorosilane (tBDMS) derivatives were formed by addition of 15 μL of 2% (w/v) methoxylamine hydrochloride (MP Biomedicals, Solon, OH) in pyridine and incubated at 45°C for 60 min. Samples were then silylated by addition of 15 μL of *N*-*tert*-butyldimethylsily-*N*-methyltrifluoroacetamide (MTBSTFA) with 1% tBDMS (Regis Technologies, Morton Grove, IL) and incubated at 45°C for 30 min.

Derivatized polar samples were injected into a GC-MS using a DB-35MS column (30 m by 0.25 mm i.d. by 0.25 μm; Agilent J&W Scientific, Santa Clara, CA) installed in an Agilent 7890B GC system integrated with an Agilent 5977a MS. Samples were injected at a GC oven temperature of 100°C which was held for 1 min before ramping to 255°C at 3.5°C/min and then to 320°C at 15°C/min and held for 3 min. Electron impact ionization was performed with the MS scanning over the range of 100 to 650 mass/charge ratio (*m/z*) for polar metabolites. Metabolite levels and mass isotopomer distributions were analyzed with an in-house MATLAB script which integrated the metabolite fragment ions and corrected for natural isotope abundances.

### LC-MS/MS analysis

For experiments in Figures 2 and 3, adherent cells were washed and scraped into 500μL of methanol and spiked with deuterated internal standards: Equisplash (Avanti Polar Lipids, Cat# 330731), SPB 18:0;O2 {d7] (Avanti Polar Lipids, Cat# 860658), SPB 18:1;O2 [D7] (Avanti Polar Lipids, Cat# 860657), and G_M3_ 18:1;O2/18:0 [D3] (Cayman Chemicals, item No. 39226). For experiments in Figures 4, 5, and 6, adherent cells and tumor punches were washed and scraped or homogenized in 500uL of methanol and spiked with deuterated internal standards: Ultimate splash (Avanti Polar Lipids, Cat#330820), SPB 18:0;O2 [D7] (Avanti Polar Lipids, Cat# 860658), SPB 18:1;O2 [D7] (Avanti Polar Lipids, Cat# 860657), and G_M3_-d_3_ 18:1;O2/18:0 [D3] (Cayman Chemicals, item No. 39226) Ten percent of the lysate was transferred to 96 well plate, dried and redissolved in 20 μL of M-PER buffer (Thermo Fisher Scientific Cat. No. 78501) for protein estimation with BCA assay. The remaining methanol extract and the tubes were vortexed and spun at 21000 × *g* for 15 minutes. The supernatants were transferred to a new tube and dried under nitrogen gas. The dried samples were resuspended in 60 μL of the initial running buffer and then analyzed with LC-MS/MS. 5 μL of sample were injected.

#### Polar lipids

Chromatographic separation and lipid species identification was performed using Q Exactive orbitrap mass spectrometer with a Vanquish Flex Binary UHPLC system (Thermo Fisher Scientific) equipped with an Kinetex C18 column, 100 × 2.1 mm, 1.7 µm particle (Phenomenex) column at 35 °C. Chromatography was performed using a gradient of 98:2 v/v water: methanol with 5 mM ammonium acetate (mobile phase A) and 50:50 v/v methanol: isopropanol with 5 mM ammonium acetate (mobile phase B), both at a flow rate of 0.2 mL/min. The liquid chromatography gradient ran with the following profile: 0 min, 30%B; 1 min, 30%B; 2 min, 70%B; 11 min, 95%B; 17 min, 30%B; 21.5 min, 30%B; 27 min, 30%B. Lipids were analyzed in positive mode using spray voltage 3.2 kV. Sweep gas flow was 1 arbitrary units, auxiliary gas flow 2 arbitrary units and sheath gas flow 40 arbitrary units, with a capillary temperature of 325 °C. Full mass spectrometry (scan range 200–2,000 m/z) was used at 140,000 resolution with 10E6 automatic gain control and a maximum injection time of 100 ms. Data dependent MS2 (Top 12) mode at 17,500 resolution with automatic gain control set at 10E5 with a maximum injection time of 50 ms was used.

#### Neutral lipids

Chromatographic separation and lipid species identification for neutral lipids was performed using Q Exactive orbitrap mass spectrometer with a Vanquish Flex Binary UHPLC system (Thermo Scientific) equipped with an Accucore C30, 150 × 2.1 mm, 2.6 µm particle (Thermo) column at 40 °C. Chromatography was performed using a gradient of 40:60 v/v water: acetonitrile with 10 mM ammonium formate and 0.1% formic acid (mobile phase A) and 10:90 v/v acetonitrile: propan-2-ol with 10 mM ammonium formate and 0.1% formic acid (mobile phase B), both at a flow rate of 0.2 ml min−1. The liquid chromatography gradient ran from 30% to 43% B from 3– 8 min, then from 43% to 50% B from 8-9 min, then 50–90% B from 9–18 min, then 90–99% B from 18–26 min, then held at 99% B from 26–30 min, before returning to 30% B in 6 min and held for a further 4 min. Neutral lipids were analyzed in positive mode using spray voltage 3.2 kV. Sweep gas flow was 1 arbitrary units, auxiliary gas flow was 2 arbitrary units, and sheath gas flow was 40 arbitrary units, with a capillary temperature of 325°C. Full MS (scan range, 200 to 2000 *m/z*) was used at 70,000 resolution with 10E6 automatic gain control and a maximum injection time of 100 ms. Data-dependent MS2 (Top 6) mode at 17,500 resolution with automatic gain control set at 10E5 with a maximum injection time of 50 ms was used.

Data were analyzed using EI-Maven. Lipid species–specific fragments used for identification and quantitation are presented in table S1. Relative abundance was calculated by normalizing to internal standards in specific to the lipid class and protein levels. Absolute abundances were calculated by normalizing to specific lipid species in Ultimate splash or specific deuterated standards added during extraction. Mass isotopologue distributions were analyzed with an in-house MATLAB script which integrated the metabolite fragment ions and corrected for natural isotope abundances.

### 13C Compound Lipid Metabolic Flux Analysis

ISA and MFA were performed to estimate the percent of newly synthesized compound lipids as well as the contribution of the tracer of interest to lipogenic acetyl-CoA pools, glycerol-3P turnover, serine pools for LCB synthesis, and the metabolic flux necessary to maintain compound lipid pools. Serine and glycerol-3P were provided in excess with unused metabolites diverted to “sink”. This allowed us to measure relative fluxes to maintain the compound lipid pools. For experiments in Figures 2-5, compound lipid pools were set to 100, without restricting any intermediates. For the experiment in Figure 6, each compound lipid pool was entered into the model as a measured pool to capture the nanomolar flux through enzymes to sustain the entered lipidome. A χ^2^ statistical test was applied to assess the goodness-of-fit using α of 0.01. Parameters for contribution of ^13^C tracers to lipogenic acetyl-CoA (*D* value) and percentage of newly synthesized compound lipids [*g*(*t*) value] and their 99% confidence intervals are then calculated using best-fit model after estimating 50 times using random initial guesses for all fluxes in the network in INCA MFA (isotopomer network compartmental analysis metabolic flux analysis) software^37,43^.

Experimental compound lipid labeling from [U-^13^C_6_]glucose and [U-^13^C_3_]serine after a 24-hour trace, as indicated in figure legends, was compared to simulated labeling using reaction networks for SM 18:1;O2/24:0, glycerophospholipids, or a broader range of compound lipids as desired and described for the experiment. Experimental replicates were entered as different wells. Estimated flux values and parameter continuation were performed with all replicates active for each of the isotopically labeled substrate used in parallel. Isotopologue distributions were entered into INCA after natural isotope correction using an in-house MATLAB script. When background noise in the isotopologue distributions were identified from ^12^C labeling wells or in regions with unexpected labeling (i.e., M+22 in a PE cultured with [U-^13^C_3_]serine, that isotopologue value was replaced with “NaN” to ignore the value during flux estimation and parameter continuation. This resulted in less than 5% of all isotopologues being removed. This was necessary due to coelution of lipid species with similar masses and the added isotopic complexity resulting from the addition of stable isotope tracing.

MFA data are plotted as 99% confidence interval. “**” indicates statistical significance by non-overlapping confidence intervals. Reaction model networks can be found in tables 1-14, with the MS1 ions used for identification and MS2 ions used for metabolite confirmation found in the table in S1. Matlab files used to generate Tables 1-14 can be found in files S1-14. Further details on assumptions can be found in Supplementary Methods.

^13^C metabolic flux analysis was conducted under the following assumptions:

1. Cells were assumed to be at metabolic steady state
2. Acetyl-CoA, glycerol-3P, and serine, as lipid precursors, are assumed to be at isotopic steady state as they turnover much faster than compound lipids, whose time-dependent fractional synthesis is modeled as the g(t) parameter. We demonstrate that glycerol-3P from [U-^13^C_6_]glucose and the serine pool from [U-^13^C_3_]serine reach steady at the beginning of the culture (**Figure S2F-G)**. This assumption has been routinely used to model acetyl-CoA contributions to fatty acid and cholesterol synthesis.
3. Cells proliferate exponentially.

### Quantification and Statistical Analysis

Data are presented as mean ± standard error of mean (SEM) of at least three biological replicates as indicated in figure legends. Statistical analysis was performed with GraphPad Prism 10.3.1 using two-tailed independent t-test to compare two groups, one-way ANOVA with Fisher’s least significant difference (LSD) post hoc test to compare more than two groups, two-way ANOVA with Fisher’s LSD post hoc test to compare two-factor study designs. For all tests, * p< 0.05, **p < 0.01, *** p< 0.001, or **** p<0.0001.

**Data S1. Unprocessed data underlying display items in the manuscript, related to Figures 1-6 and S1-6.**

**Data S2. INCA model files for all experimental conditions presented with the fractional labeling used for modeling.**

## Supplementary Figure legends

**Figure S1.**
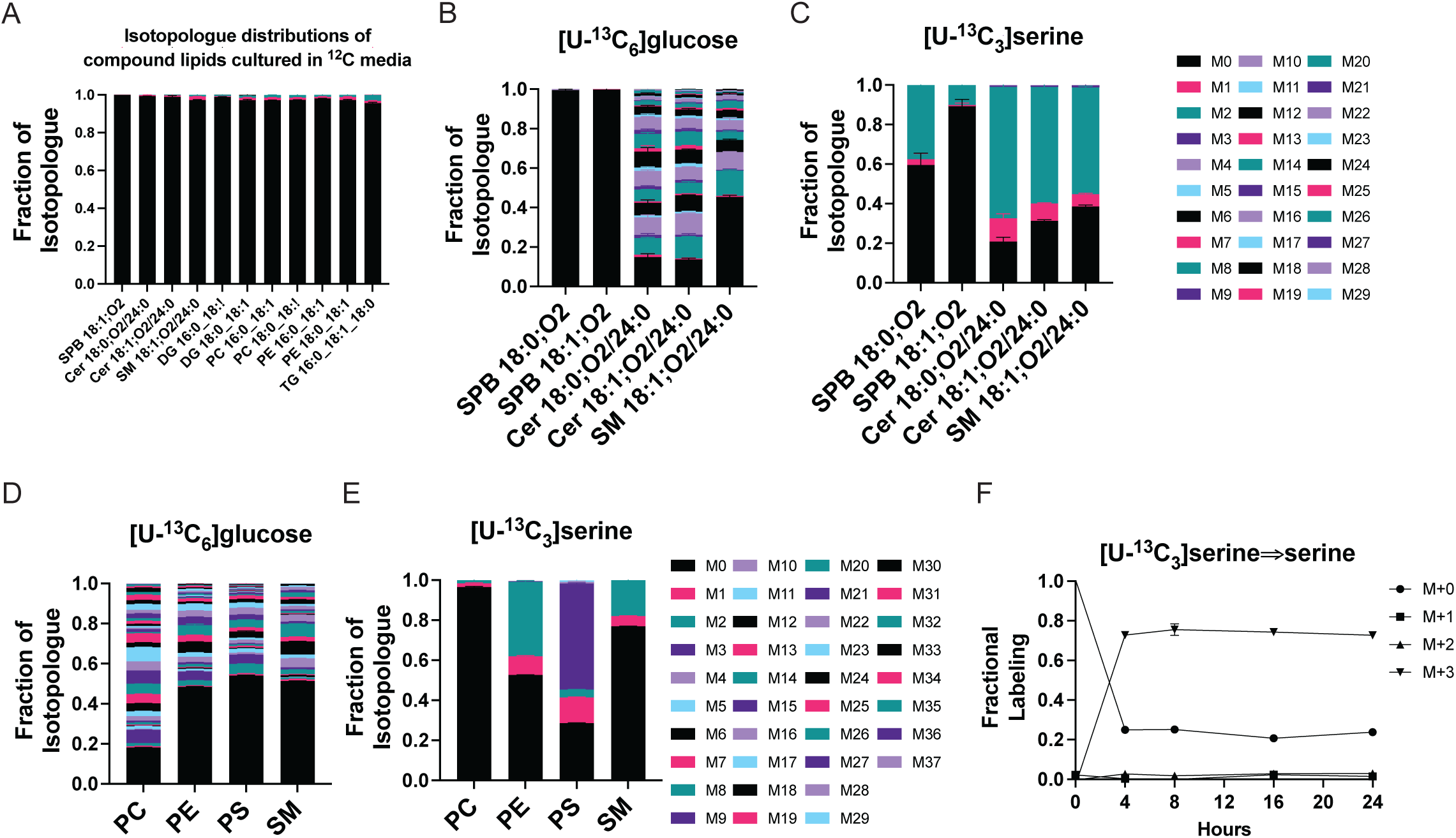
Design, implementation, and assumptions. (A) Fractional labeling for compound lipids cultured in ^12^C media showing minimal background labeling from instrument or coeluting species (n=3). (B) Fractional labeling for lipids in SM 18:1;O2/24:0 reaction network from [U-^13^C_6_]glucose (n=3). (C) Fractional labeling for lipids in SM 18:1;O2/24:0 reaction network from [U-^13^C_3_]serine (n=3). (D) Fractional labeling for compound lipids with a palmitate n-acyl chain from [U-^13^C_6_]glucose (n=3). (E) Fractional labeling for compound lipids with a palmitate n-acyl chain from [U-^13^C_6_]serine (n=3). (F) Fractional labeling of serine in A549s cultured in 20%FBS from a [U-^13^C_3_]serine tracing experiment (n=3).

**Figure S2.**
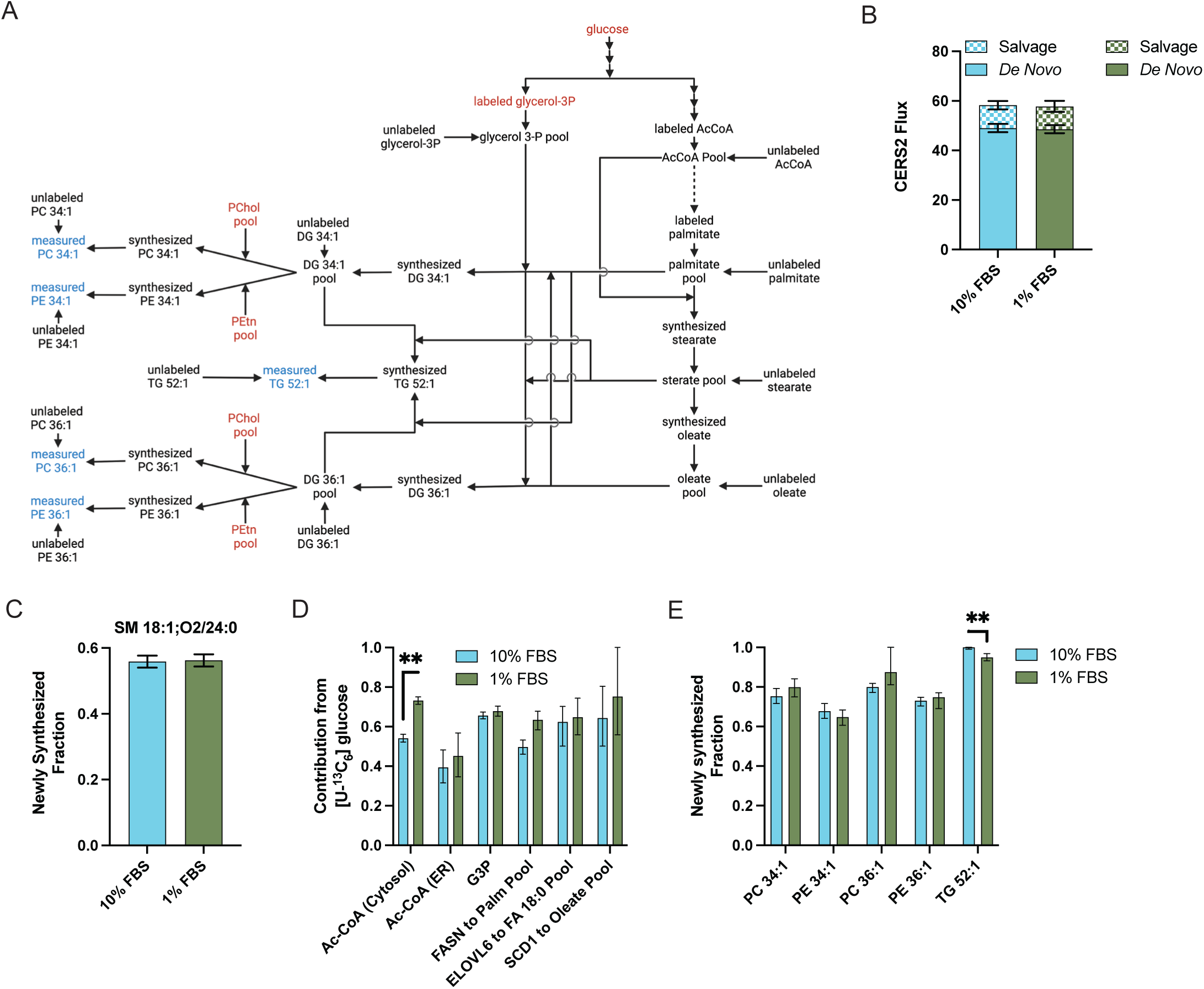
Lipid availability strongly impact acetyl-CoA and FASN flux in tissue culture. (A) Schematic of a glycerolipid MFA network. (Created with Biorender.com) (B) Comparison of CERS2 flux separated by the LCB used to generate the product of DHCer or Cer depending on sphinganine or sphingosine being the LCB used, respectively. (C) Newly synthesized fraction (g(t)) of SM 18:1;O2/24:0 after 24 hours of culture. (D) Percent contribution of [U-^13^C_6_]glucose to the different acetyl-CoA, glycerol-3P, and fatty acid pools. (E) Newly synthesized fraction (g(t)) of glycerolipids after 24 hours of culture in 1%FBS or 10%FBS. Relative abundance is calculated by normalizing to internal standard specific to lipid class. In Figure S2D-E, data shown are ratio of newly synthesized metabolite divided by the total active pool of metabolite as estimated by ^13^C MFA with 99% confidence intervals. **p < 0.01.

**Figure S3.**
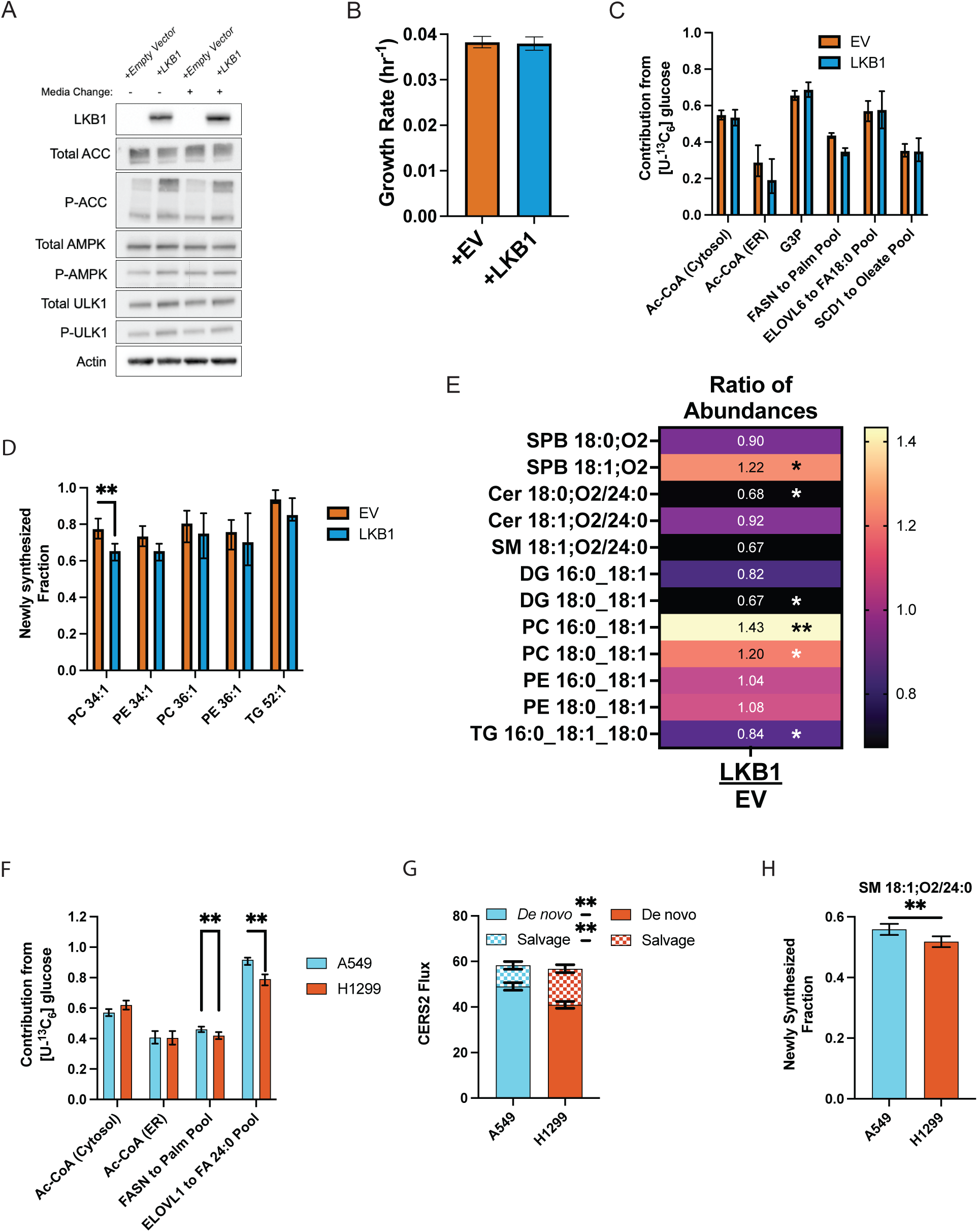
LKB1 regulates long-chain base recycling through the lysosome. (A) Immunoblot of LKB1, phosphorylated ACC (S79), ACC, phosphorylated AMPK (T172), phosphorylated ULK1 (S555), total ULK1, and β-actin on lysates from A549 cells with addition of an empty vector or LKB1. Validation of the LKB1 rescue. (B) Maximal exponential growth rate (hr^-1^) A549 +EV and +LKB1 cells (n=3). (C) Percent contribution of [U-^13^C_6_]glucose to the different acetyl-CoA and glycerol-3P pools and enzymatic flux from isotopically labeled atoms to maintain the fatty acid pool from A549 +EV and +LKB1 cells. (D) Newly synthesized fraction (g(t)) of glycerolipids after 24 hours of culture from A549 +EV and +LKB1 cells. (E) Heatmap showing ratio (+LKB1:+EV) of abundances between A549 +EV and +LKB1 cells (n=3). (F) Percent contribution of [U-^13^C_6_]glucose to the different acetyl-CoA pools or enzymatic flux from isotopically labeled atoms to maintain the fatty acid pool in A549 cells and H1299 cells. (G) Comparison of CERS2 flux separated by the LCB used to generate the product of DHCer or Cer depending on sphinganine or sphingosine being the LCB used, respectively, in A549 cells and H1299 cells. (H) Newly synthesized fraction (g(t)) of SM 18:1;O2/24:0 after 24 hours of culture in 10% FBS for A549 cells and H1299 cells. Relative abundance is calculated by normalizing to internal standard specific to lipid class. In Figures S3C-D and S3F, S3H, data shown are ratio of newly synthesized metabolite divided by the total active pool of metabolite as estimated by ^13^C MFA with 99% confidence intervals. In Figure S3G, data shown are estimated fluxes through CERS2 using sphinganine or sphingosine, simplified as *de novo* and salvage LCBs respectively, with 99% confidence intervals. ** p<0.01 for non-overlapping 99% confidence intervals.

**Figure S4.**
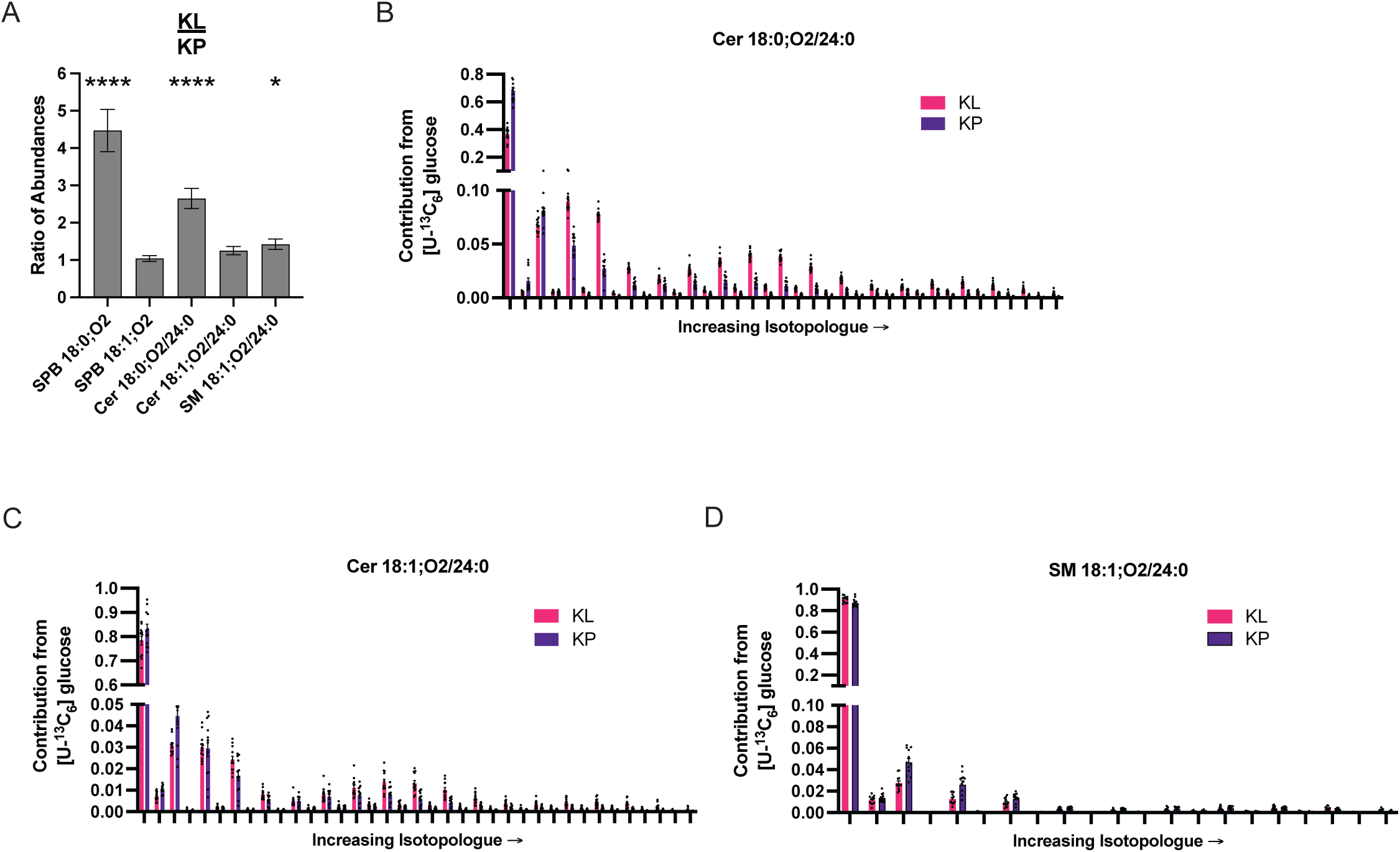
Application of CL-MFA to precision-cut lung slice culture. (A) Relative abundance for LCB from precision-cut lung slice tumors harboring KL and KP mutations (n=12). (B) Isotopologue distribution from [U-^13^C_6_]glucose for Cer 18:0;O2/24:0 in tumors after 24 hours (n=12). (C) Isotopologue distribution from [U-^13^C_6_]glucose for Cer 18:1;O2/24:0 in tumors after 24 hours (n=12). (D) Isotopologue distribution from [U-^13^C_6_]glucose for SM 18:1;O2/24:0 in tumors after 24 hours (n=12). Relative abundance is calculated by normalizing to internal standard specific to lipid class. * p<0.05, *** p<0.001, ****p<0.0001.

**Figure S5.**
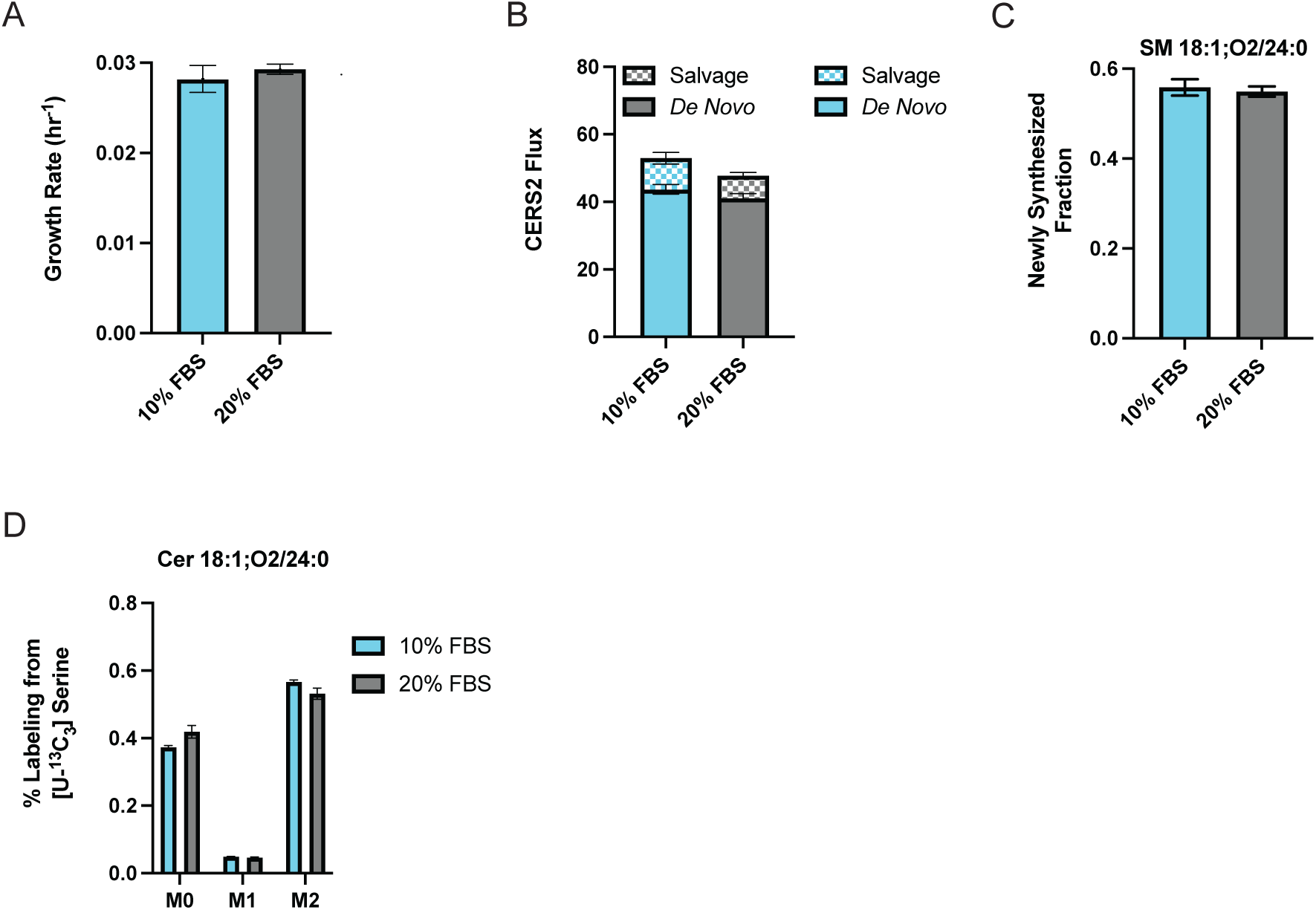
Mammalian tissue culture is inherently deficient in lipids. (A) Maximal exponential growth rate (hr^-1^) A549 cells cultured in 10% FBS or 20% FBS (n=3). (B) Comparison of CERS2 flux separated by the LCB used to generate the product of DHCer or Cer depending on sphinganine or sphingosine being the LCB used, respectively, in A549 cells cultured in 10% FBS or 20% FBS. (C) Newly synthesized fraction (g(t)) of SM 18:1;O2/24:0 after 24 hours of culture in 10% FBS for A549 cells cultured in 10% FBS or 20% FBS. (D) Cer 18:1;O2/24:0 labeling from [U-^13^C_3_]serine cultured in 10% FBS or 20% FBS for 24 hours (n=3).

**Figure S6.**
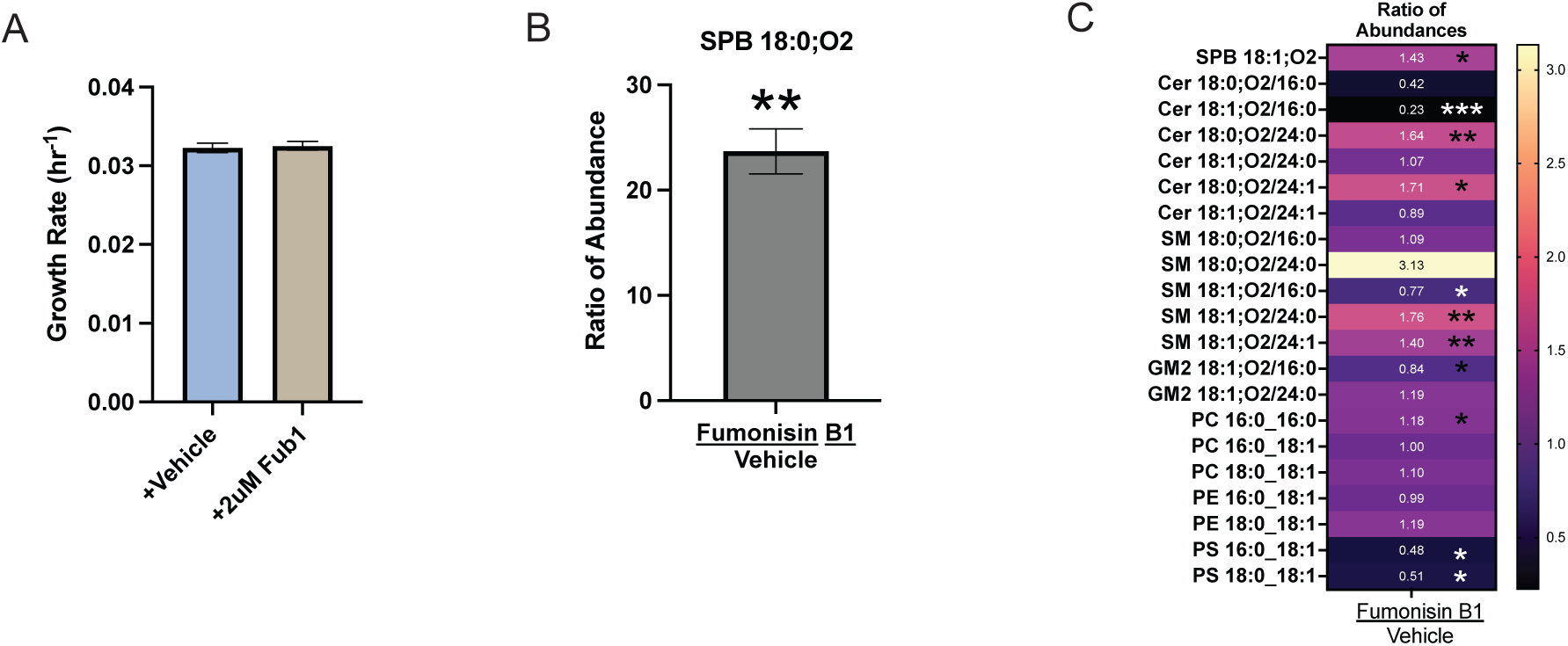
Application of CL-MFA to precision-cut lung slice culture. (A) Maximal exponential growth rate (hr^-1^) A549 cells cultured in 20% FBS with vehicle or 2μM fumonisin B1 (n=3). (B) Ratio (fumonisin B1:vehicle) of abundance of SPB 18:0;O2 in cells treated with fumonisin B1 or vehicle (n=3). (C) Heatmap showing ratio (fumonisin B1:vehicle) of abundances between A549 cells cultured in 20% FBS treated with 2μM fumonisin B1 or vehicle (n=3).

## Notes

### Competing Interest Statement

The authors have declared no competing interest.

### Summary of Updates

Small edits were made to introduction, discussion, and clarifications were added to results section. Supplemental tables and figures associated with the edited sections were updated and standardized in size and formatting to keep figures uniform and to use most updated nomenclature for lipids. Author added due to contribution analysis of samples from cell culture.

