## Supplementary material for "Modeling compound lipid homeostasis using stable isotope tracing": MFA results: A549 SL 1% FBS

### A549 1% FBS

SSR: 753.5 [627.6 774.2]

| Rxn # | Atom Transition | Flux (nanomoles) | Lower Bound | Upper Bound |
|---|---|---|---|---|
| R1 | Ac.l -> Ac.f | 438.69 | 420.89 | 456.63 |
| R2 | Ac.d -> Ac.f | 125.52 | 113.62 | 138.87 |
| R3 | Ac.f -> Ac | 564.21 | 540.55 | 588.51 |
| R4 | Ac -> AcER.l | 133.87 | 124.69 | 143.67 |
| R5 | AcER.l -> AcER.f | 133.87 | 124.69 | 143.67 |
| R6 | AcER.d -> AcER.f | 47.49 | 38.36 | 56.77 |
| R7 | AcER.f -> AcER | 181.36 | 174.68 | 189.33 |
| R8 | Ac + Ac + Ac + Ac + Ac + Ac + Ac + Ac -> Palm.s | 53.79 | 51.45 | 56.17 |
| R9 | Palm.s -> Palm | 53.79 | 51.45 | 56.17 |
| R10 | Palm.d -> Palm | 36.65 | 34.20 | 39.20 |
| R11 | Palm + AcER + AcER + AcER + AcER -> C24FA.s | 45.34 | 43.67 | 47.33 |
| R12 | C24FA.s -> C24FA | 45.34 | 43.67 | 47.33 |
| R13 | C24FA.d -> C24FA | 7.62 | 4.80 | 9.12 |
| R14 | C24FA.r -> C24FA | 4.59 | 3.43 | 5.83 |
| R15 | SerM1.l -> Ser.f | 18.85 | 17.75 | 19.94 |
| R16 | SerM2.l -> Ser.f | 5.98 | 3.85 | 8.05 |
| R17 | SerM3.l -> Ser.f | 108.69 | 106.17 | 111.31 |
| R18 | Ser.d -> Ser.f | 16.48 | 14.46 | 18.45 |
| R19 | Ser.f -> Ser | 150.00 | 150.00 | 150.00 |
| R20 | Ser -> Ser.snk | 104.90 | 103.34 | 106.45 |
| R21 | Palm + Ser -> Spha18.s + CO2 | 45.10 | 43.55 | 46.66 |
| R22 | Spha18.s -> Spha18 | 45.10 | 43.55 | 46.66 |
| R23 | Spha18.d -> Spha18 | 3.51 | 1.89 | 4.46 |
| R24 | Spha18 + C24FA -> DHCer1824.s | 48.62 | 46.56 | 50.25 |
| R25 | DHCer1824.s -> DHCer1824 | 48.62 | 46.56 | 50.25 |
| R26 | DHCer1824.d -> DHCer1824 | 0.00 | 0.00 | 1.33 |
| R27 | DHCer1824 -> Cer1824.s | 48.62 | 46.98 | 50.25 |
| R28 | Spho18 + C24FA -> Cer1824.s | 8.93 | 6.74 | 11.22 |
| R29 | Cer1824.s -> Cer1824 | 57.55 | 55.18 | 59.99 |
| R30 | Cer1824.d -> Cer1824 | 1.15 | 0.00 | 2.43 |
| R31 | Spho18.s -> Spho18 | 4.59 | 3.43 | 5.83 |
| R32 | Spho18.d -> Spho18 | 4.34 | 3.28 | 5.46 |
| R33 | Cer1824 + Chol -> SM42_1.s | 58.70 | 56.77 | 60.61 |
| R34 | SM42_1.s -> SM42_1 | 58.70 | 56.77 | 60.61 |
| R35 | SM42_1.d -> SM42_1 | 45.89 | 43.98 | 47.84 |
| R36 | SM42_1 -> Spho18.s + C24FA.r + Chol.snk | 4.59 | 3.43 | 5.83 |
| R37 | SM42_1 -> SM42_1.m | 100.00 | 100.00 | 100.00 |
