## Supplementary material for "Modeling compound lipid homeostasis using stable isotope tracing": MFA results: A549 SL 10% FBS

### A549 10% FBS

SSR: 779.6 [661.8 812.0]

| Rxn# | Atom Transition | Flux (nanomoles) | Lower Bound | Upper Bound |
|---|---|---|---|---|
| R1 | Ac.l -> Ac.f | 273.22 | 261.63 | 285.00 |
| R2 | Ac.d -> Ac.f | 205.75 | 189.80 | 222.94 |
| R3 | Ac.f -> Ac | 478.97 | 457.09 | 501.56 |
| R4 | Ac -> AcER.l | 139.07 | 128.34 | 150.46 |
| R5 | AcER.l -> AcER.f | 139.07 | 128.34 | 150.46 |
| R6 | AcER.d -> AcER.f | 55.22 | 43.77 | 66.37 |
| R7 | AcER.f -> AcER | 194.29 | 187.45 | 200.85 |
| R8 | Ac + Ac + Ac + Ac + Ac + Ac + Ac + Ac -> Palm.s | 42.49 | 40.47 | 44.56 |
| R9 | Palm.s -> Palm | 42.49 | 40.47 | 44.56 |
| R10 | Palm.d -> Palm | 49.74 | 47.42 | 52.05 |
| R11 | Palm + AcER + AcER + AcER + AcER -> C24FA.s | 48.57 | 46.86 | 50.21 |
| R12 | C24FA.s -> C24FA | 48.57 | 46.86 | 50.21 |
| R13 | C24FA.d -> C24FA | 0.00 | 0.00 | 1.83 |
| R14 | C24FA.r -> C24FA | 4.35 | 3.47 | 5.29 |
| R15 | SerM1.l -> Ser.f | 18.45 | 17.10 | 19.79 |
| R16 | SerM2.l -> Ser.f | 6.92 | 4.79 | 8.99 |
| R17 | SerM3.l -> Ser.f | 103.81 | 101.42 | 106.33 |
| R18 | Ser.d -> Ser.f | 20.81 | 18.95 | 22.47 |
| R19 | Ser.f -> Ser | 150.00 | 150.00 | 150.00 |
| R20 | Ser -> Ser.snk | 106.35 | 104.99 | 107.74 |
| R21 | Palm + Ser -> Spha18.s + CO2 | 43.65 | 42.26 | 45.01 |
| R22 | Spha18.s -> Spha18 | 43.65 | 42.26 | 45.01 |
| R23 | Spha18.d -> Spha18 | 0.00 | 0.00 | 1.20 |
| R24 | Spha18 + C24FA -> DHCer1824.s | 43.65 | 42.29 | 45.18 |
| R25 | DHCer1824.s -> DHCer1824 | 43.65 | 42.29 | 45.18 |
| R26 | DHCer1824.d -> DHCer1824 | 5.36 | 4.36 | 5.99 |
| R27 | DHCer1824 -> Cer1824.s | 49.02 | 47.37 | 50.69 |
| R28 | Spho18 + C24FA -> Cer1824.s | 9.27 | 7.52 | 10.98 |
| R29 | Cer1824.s -> Cer1824 | 58.29 | 56.24 | 60.34 |
| R30 | Cer1824.d -> Cer1824 | 0.00 | 0.00 | 0.30 |
| R31 | Spho18.s -> Spho18 | 4.35 | 3.47 | 5.29 |
| R32 | Spho18.d -> Spho18 | 4.92 | 3.97 | 5.84 |
| R33 | Cer1824 + Chol -> SM42_1.s | 58.29 | 56.24 | 60.35 |
| R34 | SM42_1.s -> SM42_1 | 58.29 | 56.24 | 60.35 |
| R35 | SM42_1.d -> SM42_1 | 46.07 | 44.28 | 47.87 |
| R36 | SM42_1 -> Spho18.s + C24FA.r + Chol.snk | 4.35 | 3.47 | 5.29 |
| R37 | SM42_1 -> SM42_1.m | 100.00 | 100.00 | 100.00 |
