## Supplementary material for "Modeling compound lipid homeostasis using stable isotope tracing": MFA results: A549 PL 1% FBS

### A549 1% FBS

SSR: 707.6 [650.4 799.4]

| Rxn # | Atom Transition | Flux (nanomoles) | Lower Bound | Upper Bound |
|---|---|---|---|---|
| R1 | Ac.l -> Ac.f | 2401 | 2318 | 2483 |
| R2 | Ac.d -> Ac.f | 879 | 824 | 935 |
| R3 | Ac.f -> Ac | 3280 | 3171 | 3386 |
| R4 | Ac -> AcER.l | 224 | 189 | 261 |
| R5 | AcER.l -> AcER.f | 224 | 189 | 261 |
| R6 | AcER.d -> AcER.f | 139 | 84 | 200 |
| R7 | AcER.f -> AcER | 363 | 327 | 406 |
| R8 | glyc.l -> glyc.f | 339 | 326 | 352 |
| R9 | glyc.d -> glyc.f | 161 | 148 | 174 |
| R10 | glyc.f -> glyc | 500 | 500 | 500 |
| R11 | glyc -> glyc.snk | 98 | 92 | 105 |
| R12 | Ac + Ac + Ac + Ac + Ac + Ac + Ac + Ac -> Palm.s | 382 | 369 | 395 |
| R13 | Palm.s -> Palm | 382 | 369 | 395 |
| R14 | Palm.d -> Palm | 220 | 188 | 263 |
| R15 | Palm + AcER -> Stea.s | 363 | 327 | 406 |
| R16 | Stea.s -> Stea | 363 | 327 | 406 |
| R17 | Stea.d -> Stea | 197 | 140 | 282 |
| R18 | Stea -> Olea.s | 302 | 223 | 406 |
| R19 | Olea.s -> Olea | 302 | 223 | 406 |
| R20 | Olea.d -> Olea | 100 | 0 | 176 |
| R21 | glyc + Palm + Olea -> DG34.s | 145 | 141 | 243 |
| R22 | DG34.s -> DG34 | 145 | 141 | 243 |
| R23 | DG34.d -> DG34 | 2 | 0 | 6 |
| R24 | DG34 + Chol -> PC34.s | 81 | 78 | 84 |
| R25 | PC34.s -> PC34 | 81 | 78 | 84 |
| R26 | PC34.d -> PC34 | 19 | 16 | 22 |
| R27 | PC34 -> PC34.m | 100 | 100 | 100 |
| R28 | DG34 + EtA -> PE34.s | 66 | 63 | 68 |
| R29 | PE34.s -> PE34 | 66 | 63 | 68 |
| R30 | PE34.d -> PE34 | 34 | 32 | 37 |
| R31 | PE34 -> PE34.m | 100 | 100 | 100 |
| R32 | glyc + Stea + Olea -> DG36.s | 257 | 160 | 261 |
| R33 | DG36.s -> DG36 | 257 | 160 | 261 |
| R34 | DG36.d -> DG36 | 0 | 0 | 8 |
| R35 | DG36 + Chol -> PC36.s | 87 | 85 | 100 |
| R36 | PC36.s -> PC36 | 87 | 85 | 100 |
| R37 | PC36.d -> PC36 | 13 | 0 | 15 |

| | | | | |
|---|---|---|---|---|
| R38 | PC36 -> PC36.m | 100 | 100 | 100 |
| R39 | DG36 + EtA -> PE36.s | 75 | 73 | 77 |
| R40 | PE36.s -> PE36 | 75 | 73 | 77 |
| R41 | PE36.d -> PE36 | 25 | 23 | 27 |
| R42 | PE36 -> PE36.m | 100 | 100 | 100 |
| R43 | DG34 + Stea -> TG52_1.s | 0 | 0 | 97 |
| R44 | DG36 + Palm -> TG52_1.s | 95 | 0 | 97 |
| R45 | TG52_1.s -> TG52_1 | 95 | 93 | 97 |
| R46 | TG52_1.d -> TG52_1 | 5 | 3 | 7 |
| R47 | TG52_1 -> TG52_1.m | 100 | 100 | 100 |
