## Supplementary material for "Modeling compound lipid homeostasis using stable isotope tracing": MFA results: A549 PL 10% FBS

### A549 10% FBS

SSR: 770.7 [650.4 799.4]

| Rxn# | Atom Transition | Flux (nanomoles) | Lower Bound | Upper Bound |
|---|---|---|---|---|
| R1 | Ac.l -> Ac.f | 1331 | 1286 | 1376 |
| R2 | Ac.d -> Ac.f | 1127 | 1077 | 1176 |
| R3 | Ac.f -> Ac | 2458 | 2383 | 2530 |
| R4 | Ac -> AcER.l | 230 | 207 | 254 |
| R5 | AcER.l -> AcER.f | 230 | 207 | 254 |
| R6 | AcER.d -> AcER.f | 87 | 42 | 135 |
| R7 | AcER.f -> AcER | 317 | 284 | 354 |
| R8 | glyc.l -> glyc.f | 328 | 320 | 337 |
| R9 | glyc.d -> glyc.f | 172 | 163 | 180 |
| R10 | glyc.f -> glyc | 500 | 500 | 500 |
| R11 | glyc -> glyc.snk | 104 | 99 | 109 |
| R12 | Ac + Ac + Ac + Ac + Ac + Ac + Ac + Ac -> Palm.s | 278 | 269 | 288 |
| R13 | Palm.s -> Palm | 278 | 269 | 288 |
| R14 | Palm.d -> Palm | 281 | 253 | 315 |
| R15 | Palm + AcER -> Stea.s | 317 | 284 | 354 |
| R16 | Stea.s -> Stea | 317 | 284 | 354 |
| R17 | Stea.d -> Stea | 191 | 150 | 241 |
| R18 | Stea -> Olea.s | 255 | 198 | 319 |
| R19 | Olea.s -> Olea | 255 | 198 | 319 |
| R20 | Olea.d -> Olea | 141 | 78 | 197 |
| R21 | glyc + Palm + Olea -> DG34.s | 143 | 140 | 163 |
| R22 | DG34.s -> DG34 | 143 | 140 | 163 |
| R23 | DG34.d -> DG34 | 8 | 6 | 12 |
| R24 | DG34 + Chol -> PC34.s | 80 | 78 | 82 |
| R25 | PC34.s -> PC34 | 80 | 78 | 82 |
| R26 | PC34.d -> PC34 | 20 | 18 | 22 |
| R27 | PC34 -> PC34.m | 100 | 100 | 100 |
| R28 | DG34 + EtA -> PE34.s | 72 | 69 | 74 |
| R29 | PE34.s -> PE34 | 72 | 69 | 74 |
| R30 | PE34.d -> PE34 | 28 | 26 | 31 |
| R31 | PE34 -> PE34.m | 100 | 100 | 100 |
| R32 | glyc + Stea + Olea -> DG36.s | 253 | 231 | 256 |
| R33 | DG36.s -> DG36 | 253 | 231 | 256 |
| R34 | DG36.d -> DG36 | 0 | 0 | 3 |
| R35 | DG36 + Chol -> PC36.s | 80 | 78 | 82 |
| R36 | PC36.s -> PC36 | 80 | 78 | 82 |
| R37 | PC36.d -> PC36 | 20 | 18 | 22 |

| | | | | |
|---|---|---|---|---|
| R38 | PC36 -> PC36.m | 100 | 100 | 100 |
| R39 | DG36 + EtA -> PE36.s | 73 | 71 | 75 |
| R40 | PE36.s -> PE36 | 73 | 71 | 75 |
| R41 | PE36.d -> PE36 | 27 | 25 | 29 |
| R42 | PE36 -> PE36.m | 100 | 100 | 100 |
| R43 | DG34 + Stea -> TG52_1.s | 0 | 0 | 21 |
| R44 | DG36 + Palm -> TG52_1.s | 100 | 79 | NaN |
| R45 | TG52_1.s -> TG52_1 | 100 | 99 | 100 |
| R46 | TG52_1.d -> TG52_1 | 0 | 0 | 1 |
| R47 | TG52_1 -> TG52_1.m | 100 | 100 | 100 |
