## Supplementary material for "Modeling compound lipid homeostasis using stable isotope tracing": MFA results: A549 SL +EV 10% FBS

### A549 +EV 10% FBS

SSR: 837.0 [709.3 864.5]

| Rxn# | Atom Transition | Flux (nanomoles) | Lower Bound | Upper Bound |
|---|---|---|---|---|
| R1 | Ac.l -> Ac.f | 286.13 | 277.63 | 294.69 |
| R2 | Ac.d -> Ac.f | 208.92 | 198.24 | 219.99 |
| R3 | Ac.f -> Ac | 495.05 | 481.20 | 508.99 |
| R4 | Ac -> AcER.l | 160.10 | 153.85 | 166.53 |
| R5 | AcER.l -> AcER.f | 160.10 | 153.85 | 166.53 |
| R6 | AcER.d -> AcER.f | 68.31 | 61.23 | 75.37 |
| R7 | AcER.f -> AcER | 228.41 | 223.90 | 232.91 |
| R8 | Ac + Ac + Ac + Ac + Ac + Ac + Ac + Ac -> Palm.s | 41.87 | 40.38 | 43.37 |
| R9 | Palm.s -> Palm | 41.87 | 40.38 | 43.37 |
| R10 | Palm.d -> Palm | 77.17 | 75.22 | 79.12 |
| R11 | Palm + AcER + AcER + AcER + AcER -> C24FA.s | 57.10 | 55.97 | 58.23 |
| R12 | C24FA.s -> C24FA | 57.10 | 55.97 | 58.23 |
| R13 | C24FA.d -> C24FA | 13.60 | 12.43 | 14.58 |
| R14 | C24FA.r -> C24FA | 1.62 | 1.37 | 1.89 |
| R15 | SerM1.l -> Ser.f | 10.19 | 9.58 | 10.80 |
| R16 | SerM2.l -> Ser.f | 3.54 | 2.14 | 4.90 |
| R17 | SerM3.l -> Ser.f | 65.40 | 63.95 | 66.90 |
| R18 | Ser.d -> Ser.f | 20.87 | 20.05 | 21.66 |
| R19 | Ser.f -> Ser | 100.00 | 100.00 | 100.00 |
| R20 | Ser -> Ser.snk | 38.06 | 36.84 | 39.28 |
| R21 | Palm + Ser -> Spha18.s + CO2 | 61.94 | 60.72 | 63.16 |
| R22 | Spha18.s -> Spha18 | 61.94 | 60.72 | 63.16 |
| R23 | Spha18.d -> Spha18 | 0.00 | 0.00 | 0.39 |
| R24 | Spha18 + C24FA -> DHCer1824.s | 61.94 | 60.72 | 63.16 |
| R25 | DHCer1824.s -> DHCer1824 | 61.94 | 60.72 | 63.16 |
| R26 | DHCer1824.d -> DHCer1824 | 0.00 | 0.00 | 0.55 |
| R27 | DHCer1824 -> Cer1824.s | 61.94 | 60.72 | 63.16 |
| R28 | Spho18 + C24FA -> Cer1824.s | 10.38 | 9.35 | 11.42 |
| R29 | Cer1824.s -> Cer1824 | 72.32 | 71.04 | 73.59 |
| R30 | Cer1824.d -> Cer1824 | 0.00 | 0.00 | 0.30 |
| R31 | Spho18.s -> Spho18 | 1.62 | 1.37 | 1.89 |
| R32 | Spho18.d -> Spho18 | 8.77 | 7.90 | 9.63 |
| R33 | Cer1824 + Chol -> SM42_1.s | 72.32 | 71.04 | 73.59 |
| R34 | SM42_1.s -> SM42_1 | 72.32 | 71.04 | 73.59 |
| R35 | SM42_1.d -> SM42_1 | 29.30 | 28.07 | 30.53 |
| R36 | SM42_1 -> Spho18.s + C24FA.r + Chol.snk | 1.62 | 1.37 | 1.89 |
| R37 | SM42_1 -> SM42_1.m | 100.00 | 100.00 | 100.00 |
