## Supplementary material for "Modeling compound lipid homeostasis using stable isotope tracing": MFA results: A549 SL +LKB1 10% FBS

### A549 +LKB1 10% FBS

SSR: 851.0 [755.9 915.9]

| Rxn# | Atom Transition | Flux (nanomoles) | Lower Bound | Upper Bound |
|---|---|---|---|---|
| R1 | Ac.l -> Ac.f | 250.30 | 242.53 | 258.12 |
| R2 | Ac.d -> Ac.f | 198.38 | 187.08 | 210.19 |
| R3 | Ac.f -> Ac | 448.68 | 434.98 | 462.57 |
| R4 | Ac -> AcER.l | 158.31 | 152.09 | 164.77 |
| R5 | AcER.l -> AcER.f | 158.31 | 152.09 | 164.77 |
| R6 | AcER.d -> AcER.f | 68.88 | 62.09 | 75.60 |
| R7 | AcER.f -> AcER | 227.19 | 222.54 | 231.85 |
| R8 | Ac + Ac + Ac + Ac + Ac + Ac + Ac + Ac -> Palm.s | 36.30 | 34.87 | 37.73 |
| R9 | Palm.s -> Palm | 36.30 | 34.87 | 37.73 |
| R10 | Palm.d -> Palm | 78.66 | 76.74 | 80.59 |
| R11 | Palm + AcER + AcER + AcER + AcER -> C24FA.s | 56.80 | 55.63 | 57.96 |
| R12 | C24FA.s -> C24FA | 56.80 | 55.63 | 57.96 |
| R13 | C24FA.d -> C24FA | 15.58 | 13.60 | 17.59 |
| R14 | C24FA.r -> C24FA | 3.24 | 2.61 | 3.91 |
| R15 | SerM1.l -> Ser.f | 8.95 | 8.25 | 9.65 |
| R16 | SerM2.l -> Ser.f | 3.61 | 2.19 | 5.00 |
| R17 | SerM3.l -> Ser.f | 64.32 | 62.80 | 65.90 |
| R18 | Ser.d -> Ser.f | 23.11 | 21.88 | 24.33 |
| R19 | Ser.f -> Ser | 100.00 | 100.00 | 100.00 |
| R20 | Ser -> Ser.snk | 41.84 | 40.43 | 43.23 |
| R21 | Palm + Ser -> Spha18.s + CO2 | 58.16 | 56.77 | 59.57 |
| R22 | Spha18.s -> Spha18 | 58.16 | 56.77 | 59.57 |
| R23 | Spha18.d -> Spha18 | 0.00 | 0.00 | 0.59 |
| R24 | Spha18 + C24FA -> DHCer1824.s | 58.16 | 56.77 | 59.57 |
| R25 | DHCer1824.s -> DHCer1824 | 58.16 | 56.77 | 59.57 |
| R26 | DHCer1824.d -> DHCer1824 | 1.39 | 0.26 | 2.52 |
| R27 | DHCer1824 -> Cer1824.s | 59.55 | 58.35 | 60.76 |
| R28 | Spho18 + C24FA -> Cer1824.s | 17.46 | 16.00 | 18.96 |
| R29 | Cer1824.s -> Cer1824 | 77.01 | 75.57 | 78.47 |
| R30 | Cer1824.d -> Cer1824 | 0.00 | 0.00 | 0.27 |
| R31 | Spho18.s -> Spho18 | 3.24 | 2.61 | 3.91 |
| R32 | Spho18.d -> Spho18 | 14.22 | 13.07 | 15.39 |
| R33 | Cer1824 + Chol -> SM42_1.s | 77.01 | 75.57 | 78.47 |
| R34 | SM42_1.s -> SM42_1 | 77.01 | 75.57 | 78.47 |
| R35 | SM42_1.d -> SM42_1 | 26.23 | 24.95 | 27.51 |
| R36 | SM42_1 -> Spho18.s + C24FA.r + Chol.snk | 3.24 | 2.61 | 3.91 |
| R37 | SM42_1 -> SM42_1.m | 100.00 | 100.00 | 100.00 |
