## Supplementary material for "Modeling compound lipid homeostasis using stable isotope tracing": MFA results: A549 PL +EV 10% FBS

### A549 +EV 10% FBS

SSR: 690.7 [650.4 799.4]

| Rxn# | Atom Transition | Flux (nanomoles) | Lower Bound | Upper Bound |
|---|---|---|---|---|
| R1 | Ac.l -> Ac.f | 1027 | 979 | 1075 |
| R2 | Ac.d -> Ac.f | 846 | 799 | 894 |
| R3 | Ac.f -> Ac | 1873 | 1797 | 1948 |
| R4 | Ac -> AcER.l | 173 | 155 | 192 |
| R5 | AcER.l -> AcER.f | 173 | 155 | 192 |
| R6 | AcER.d -> AcER.f | 158 | 96 | 226 |
| R7 | AcER.f -> AcER | 331 | 280 | 388 |
| R8 | glyc.l -> glyc.f | 328 | 315 | 341 |
| R9 | glyc.d -> glyc.f | 172 | 159 | 185 |
| R10 | glyc.f -> glyc | 500 | 500 | 500 |
| R11 | glyc -> glyc.snk | 100 | 91 | 109 |
| R12 | Ac + Ac + Ac + Ac + Ac + Ac + Ac + Ac -> Palm.s | 213 | 203 | 222 |
| R13 | Palm.s -> Palm | 213 | 203 | 222 |
| R14 | Palm.d -> Palm | 362 | 315 | 417 |
| R15 | Palm + AcER -> Stea.s | 331 | 280 | 388 |
| R16 | Stea.s -> Stea | 331 | 280 | 388 |
| R17 | Stea.d -> Stea | 290 | 215 | 352 |
| R18 | Stea -> Olea.s | 371 | 276 | 408 |
| R19 | Olea.s -> Olea | 371 | 276 | 408 |
| R20 | Olea.d -> Olea | 29 | 0 | 123 |
| R21 | glyc + Palm + Olea -> DG34.s | 151 | 147 | 200 |
| R22 | DG34.s -> DG34 | 151 | 147 | 200 |
| R23 | DG34.d -> DG34 | 16 | 12 | 22 |
| R24 | DG34 + Chol -> PC34.s | 86 | 83 | 88 |
| R25 | PC34.s -> PC34 | 86 | 83 | 88 |
| R26 | PC34.d -> PC34 | 14 | 12 | 17 |
| R27 | PC34 -> PC34.m | 100 | 100 | 100 |
| R28 | DG34 + EtA -> PE34.s | 81 | 78 | 84 |
| R29 | PE34.s -> PE34 | 81 | 78 | 84 |
| R30 | PE34.d -> PE34 | 19 | 16 | 22 |
| R31 | PE34 -> PE34.m | 100 | 100 | 100 |
| R32 | glyc + Stea + Olea -> DG36.s | 250 | 198 | 255 |
| R33 | DG36.s -> DG36 | 250 | 198 | 255 |
| R34 | DG36.d -> DG36 | 0 | 0 | 22 |
| R35 | DG36 + Chol -> PC36.s | 80 | 78 | 87 |
| R36 | PC36.s -> PC36 | 80 | 78 | 87 |
| R37 | PC36.d -> PC36 | 20 | 13 | 22 |

| | | | | |
|---|---|---|---|---|
| R38 | PC36 -> PC36.m | 100 | 100 | 100 |
| R39 | DG36 + EtA -> PE36.s | 76 | 74 | 82 |
| R40 | PE36.s -> PE36 | 76 | 74 | 82 |
| R41 | PE36.d -> PE36 | 24 | 18 | 26 |
| R42 | PE36 -> PE36.m | 100 | 100 | 100 |
| R43 | DG34 + Stea -> TG52_1.s | 0 | 0 | 55 |
| R44 | DG36 + Palm -> TG52_1.s | 94 | 42 | 99 |
| R45 | TG52_1.s -> TG52_1 | 94 | 92 | 99 |
| R46 | TG52_1.d -> TG52_1 | 6 | 1 | 8 |
| R47 | TG52_1 -> TG52_1.m | 100 | 100 | 100 |
