## Supplementary material for "Modeling compound lipid homeostasis using stable isotope tracing": MFA results: A549 PL +LKB1 10% FBS

### A549 +LKB1 10% FBS

SSR: 704.5 [633.3 780.5]

| Rxn# | Atom Transition | Flux (nanomoles) | Lower Bound | Upper Bound |
|---|---|---|---|---|
| R1 | Ac.l -> Ac.f | 731 | 672 | 790 |
| R2 | Ac.d -> Ac.f | 638 | 580 | 697 |
| R3 | Ac.f -> Ac | 1369 | 1274 | 1462 |
| R4 | Ac -> AcER.l | 130 | 112 | 151 |
| R5 | AcER.l -> AcER.f | 130 | 112 | 151 |
| R6 | AcER.d -> AcER.f | 233 | 132 | 349 |
| R7 | AcER.f -> AcER | 364 | 274 | 470 |
| R8 | glyc.l -> glyc.f | 343 | 324 | 364 |
| R9 | glyc.d -> glyc.f | 157 | 136 | 176 |
| R10 | glyc.f -> glyc | 500 | 500 | 500 |
| R11 | glyc -> glyc.snk | 139 | 127 | 153 |
| R12 | Ac + Ac + Ac + Ac + Ac + Ac + Ac + Ac -> Palm.s | 155 | 143 | 167 |
| R13 | Palm.s -> Palm | 155 | 143 | 167 |
| R14 | Palm.d -> Palm | 425 | 339 | 529 |
| R15 | Palm + AcER -> Stea.s | 364 | 274 | 470 |
| R16 | Stea.s -> Stea | 364 | 274 | 470 |
| R17 | Stea.d -> Stea | 227 | 119 | 303 |
| R18 | Stea -> Olea.s | 361 | 245 | 373 |
| R19 | Olea.s -> Olea | 361 | 245 | 373 |
| R20 | Olea.d -> Olea | 0 | 0 | 114 |
| R21 | glyc + Palm + Olea -> DG34.s | 131 | 125 | 146 |
| R22 | DG34.s -> DG34 | 131 | 125 | 146 |
| R23 | DG34.d -> DG34 | 69 | 64 | 79 |
| R24 | DG34 + Chol -> PC34.s | 100 | 98 | 100 |
| R25 | PC34.s -> PC34 | 100 | 98 | 100 |
| R26 | PC34.d -> PC34 | 0 | 0 | 2 |
| R27 | PC34 -> PC34.m | 100 | 100 | 100 |
| R28 | DG34 + EtA -> PE34.s | 100 | 98 | 100 |
| R29 | PE34.s -> PE34 | 100 | 98 | 100 |
| R30 | PE34.d -> PE34 | 0 | 0 | 2 |
| R31 | PE34 -> PE34.m | 100 | 100 | 100 |
| R32 | glyc + Stea + Olea -> DG36.s | 230 | 211 | 238 |
| R33 | DG36.s -> DG36 | 230 | 211 | 238 |
| R34 | DG36.d -> DG36 | 0 | 0 | 35 |
| R35 | DG36 + Chol -> PC36.s | 75 | 71 | 86 |
| R36 | PC36.s -> PC36 | 75 | 71 | 86 |
| R37 | PC36.d -> PC36 | 25 | 14 | 29 |

| | | | | |
|---|---|---|---|---|
| R38 | PC36 -> PC36.m | 100 | 100 | 100 |
| R39 | DG36 + EtA -> PE36.s | 70 | 67 | 81 |
| R40 | PE36.s -> PE36 | 70 | 67 | 81 |
| R41 | PE36.d -> PE36 | 30 | 19 | 33 |
| R42 | PE36 -> PE36.m | 100 | 100 | 100 |
| R43 | DG34 + Stea -> TG52_1.s | 0 | 0 | 23 |
| R44 | DG36 + Palm -> TG52_1.s | 85 | 68 | 94 |
| R45 | TG52_1.s -> TG52_1 | 85 | 82 | 94 |
| R46 | TG52_1.d -> TG52_1 | 15 | 6 | 18 |
| R47 | TG52_1 -> TG52_1.m | 100 | 100 | 100 |
