## Supplementary material for "Modeling compound lipid homeostasis using stable isotope tracing": MFA results: H1299 SL 10% FBS

### H1299 10% FBS

SSR: 825.3 [717.8 874.0]

| Rxn# | Atom Transition | Flux (nanomoles) | Lower Bound | Upper Bound |
|---|---|---|---|---|
| R1 | Ac.l -> Ac.f | 194.90 | 185.84 | 204.07 |
| R2 | Ac.d -> Ac.f | 119.39 | 110.24 | 129.00 |
| R3 | Ac.f -> Ac | 314.29 | 299.74 | 329.00 |
| R4 | Ac -> AcER.l | 93.86 | 88.08 | 99.85 |
| R5 | AcER.l -> AcER.f | 93.86 | 88.08 | 99.85 |
| R6 | AcER.d -> AcER.f | 50.21 | 44.34 | 56.06 |
| R7 | AcER.f -> AcER | 144.07 | 138.90 | 149.15 |
| R8 | Ac + Ac + Ac + Ac + Ac + Ac + Ac + Ac -> Palm.s | 27.55 | 26.12 | 29.00 |
| R9 | Palm.s -> Palm | 27.55 | 26.12 | 29.00 |
| R10 | Palm.d -> Palm | 38.19 | 36.56 | 39.81 |
| R11 | Palm + AcER + AcER + AcER + AcER -> C24FA.s | 36.02 | 34.73 | 37.29 |
| R12 | C24FA.s -> C24FA | 36.02 | 34.73 | 37.29 |
| R13 | C24FA.d -> C24FA | 0.00 | 0.00 | 0.44 |
| R14 | C24FA.r -> C24FA | 9.54 | 8.08 | 11.16 |
| R15 | SerM1.l -> Ser.f | 22.14 | 21.30 | 22.97 |
| R16 | SerM2.l -> Ser.f | 6.67 | 5.33 | 7.98 |
| R17 | SerM3.l -> Ser.f | 54.88 | 53.48 | 56.32 |
| R18 | Ser.d -> Ser.f | 16.32 | 15.12 | 17.44 |
| R19 | Ser.f -> Ser | 100.00 | 100.00 | 100.00 |
| R20 | Ser -> Ser.snk | 70.27 | 69.30 | 71.28 |
| R21 | Palm + Ser -> Spha18.s + CO2 | 29.73 | 28.72 | 30.70 |
| R22 | Spha18.s -> Spha18 | 29.73 | 28.72 | 30.70 |
| R23 | Spha18.d -> Spha18 | 0.00 | 0.00 | 0.76 |
| R24 | Spha18 + C24FA -> DHCer1824.s | 29.73 | 28.76 | 30.72 |
| R25 | DHCer1824.s -> DHCer1824 | 29.73 | 28.76 | 30.72 |
| R26 | DHCer1824.d -> DHCer1824 | 11.22 | 10.40 | 11.99 |
| R27 | DHCer1824 -> Cer1824.s | 40.95 | 39.46 | 42.46 |
| R28 | Spho18 + C24FA -> Cer1824.s | 15.83 | 14.14 | 17.63 |
| R29 | Cer1824.s -> Cer1824 | 56.78 | 54.66 | 58.93 |
| R30 | Cer1824.d -> Cer1824 | 0.00 | 0.00 | 0.17 |
| R31 | Spho18.s -> Spho18 | 9.54 | 8.08 | 11.16 |
| R32 | Spho18.d -> Spho18 | 6.29 | 5.42 | 7.12 |
| R33 | Cer1824 + Chol -> SM42_1.s | 56.78 | 54.66 | 58.93 |
| R34 | SM42_1.s -> SM42_1 | 56.78 | 54.66 | 58.93 |
| R35 | SM42_1.d -> SM42_1 | 52.76 | 51.00 | 54.56 |
| R36 | SM42_1 -> Spho18.s + C24FA.r + Chol.snk | 9.54 | 8.08 | 11.16 |
| R37 | SM42_1 -> SM42_1.m | 100.00 | 100.00 | 100.00 |
