## Supplementary material for "Modeling compound lipid homeostasis using stable isotope tracing": MFA results: KL lung Slice SL 2% FBS

### KL Lung Slice 2% FBS

SSR: 760.5 [648.5 797.3]

| Rxn# | Atom Transition | Flux (nanomoles) | Lower Bound | Upper Bound |
|---|---|---|---|---|
| R1 | Ac.l -> Ac.f | 57.55 | 52.48 | 62.75 |
| R2 | Ac.d -> Ac.f | 12.07 | 10.11 | 14.21 |
| R3 | Ac.f -> Ac | 69.62 | 63.39 | 76.03 |
| R4 | Ac -> AcER.l | 25.75 | 23.36 | 28.24 |
| R5 | AcER.l -> AcER.f | 25.75 | 23.36 | 28.24 |
| R6 | AcER.d -> AcER.f | 15.33 | 13.69 | 17.06 |
| R7 | AcER.f -> AcER | 41.09 | 37.51 | 44.71 |
| R8 | Ac + Ac + Ac + Ac + Ac + Ac + Ac + Ac -> Palm.s | 5.48 | 4.95 | 6.03 |
| R9 | Palm.s -> Palm | 5.48 | 4.95 | 6.03 |
| R10 | Palm.d -> Palm | 21.15 | 19.45 | 22.82 |
| R11 | Palm + AcER + AcER + AcER + AcER -> C24FA.s | 10.27 | 9.38 | 11.18 |
| R12 | C24FA.s -> C24FA | 10.27 | 9.38 | 11.18 |
| R13 | C24FA.d -> C24FA | 6.08 | 4.92 | 7.30 |
| R14 | C24FA.r -> C24FA | 3.46 | 1.65 | 5.36 |
| R15 | SerM1.l -> Ser.f | 17.95 | 17.11 | 18.78 |
| R16 | SerM2.l -> Ser.f | 4.22 | 2.84 | 5.57 |
| R17 | SerM3.l -> Ser.f | 46.13 | 44.82 | 47.47 |
| R18 | Ser.d -> Ser.f | 31.70 | 30.43 | 32.99 |
| R19 | Ser.f -> Ser | 100.00 | 100.00 | 100.00 |
| R20 | Ser -> Ser.snk | 83.64 | 82.21 | 85.06 |
| R21 | Palm + Ser -> Spha18.s + CO2 | 16.36 | 14.94 | 17.79 |
| R22 | Spha18.s -> Spha18 | 16.36 | 14.94 | 17.79 |
| R23 | Spha18.d -> Spha18 | 0.00 | 0.00 | 0.45 |
| R24 | Spha18 + C24FA -> DHCer1824.s | 16.36 | 14.94 | 17.79 |
| R25 | DHCer1824.s -> DHCer1824 | 16.36 | 14.94 | 17.79 |
| R26 | DHCer1824.d -> DHCer1824 | 1.12 | 0.58 | 1.53 |
| R27 | DHCer1824 -> Cer1824.s | 17.47 | 15.97 | 18.98 |
| R28 | Spho18 + C24FA -> Cer1824.s | 3.46 | 1.65 | 5.36 |
| R29 | Cer1824.s -> Cer1824 | 20.94 | 18.95 | 23.04 |
| R30 | Cer1824.d -> Cer1824 | 51.61 | 46.44 | 57.16 |
| R31 | Spho18.s -> Spho18 | 3.46 | 1.65 | 5.36 |
| R32 | Spho18.d -> Spho18 | 0.00 | 0.00 | 0.07 |
| R33 | Cer1824 + Chol -> SM42_1.s | 72.55 | 65.95 | 79.47 |
| R34 | SM42_1.s -> SM42_1 | 72.55 | 65.95 | 79.47 |
| R35 | SM42_1.d -> SM42_1 | 30.92 | 24.27 | 37.29 |
| R36 | SM42_1 -> Spho18.s + C24FA.r + Chol.snk | 3.46 | 1.65 | 5.36 |
| R37 | SM42_1 -> SM42_1.m | 100.00 | 100.00 | 100.00 |
