## Supplementary material for "Modeling compound lipid homeostasis using stable isotope tracing": MFA results: KP lung Slice SL 2% FBS

### KP Lung Slice 2% FBS

SSR: 845.9 [707.4 862.4]

| Rxn# | Atom Transition | Flux (nanomoles) | Lower Bound | Upper Bound |
|---|---|---|---|---|
| R1 | Ac.l -> Ac.f | 33.78 | 31.16 | 36.49 |
| R2 | Ac.d -> Ac.f | 11.75 | 9.50 | 14.24 |
| R3 | Ac.f -> Ac | 45.52 | 41.79 | 49.38 |
| R4 | Ac -> AcER.l | 23.55 | 21.73 | 25.47 |
| R5 | AcER.l -> AcER.f | 23.55 | 21.73 | 25.47 |
| R6 | AcER.d -> AcER.f | 29.39 | 27.21 | 31.61 |
| R7 | AcER.f -> AcER | 52.94 | 50.02 | 55.86 |
| R8 | Ac + Ac + Ac + Ac + Ac + Ac + Ac + Ac -> Palm.s | 2.75 | 2.43 | 3.08 |
| R9 | Palm.s -> Palm | 2.75 | 2.43 | 3.08 |
| R10 | Palm.d -> Palm | 24.30 | 23.04 | 25.56 |
| R11 | Palm + AcER + AcER + AcER + AcER -> C24FA.s | 13.23 | 12.51 | 13.96 |
| R12 | C24FA.s -> C24FA | 13.23 | 12.51 | 13.96 |
| R13 | C24FA.d -> C24FA | 10.62 | 6.98 | 15.31 |
| R14 | C24FA.r -> C24FA | 15.14 | 10.55 | 20.87 |
| R15 | SerM1.l -> Ser.f | 26.74 | 25.89 | 27.59 |
| R16 | SerM2.l -> Ser.f | 6.41 | 5.38 | 7.43 |
| R17 | SerM3.l -> Ser.f | 40.57 | 39.58 | 41.58 |
| R18 | Ser.d -> Ser.f | 26.27 | 25.30 | 27.25 |
| R19 | Ser.f -> Ser | 100.00 | 100.00 | 100.00 |
| R20 | Ser -> Ser.snk | 86.19 | 85.22 | 87.14 |
| R21 | Palm + Ser -> Spha18.s + CO2 | 13.81 | 12.86 | 14.78 |
| R22 | Spha18.s -> Spha18 | 13.81 | 12.86 | 14.78 |
| R23 | Spha18.d -> Spha18 | 0.00 | 0.00 | 1.83 |
| R24 | Spha18 + C24FA -> DHCer1824.s | 13.81 | 12.86 | 15.78 |
| R25 | DHCer1824.s -> DHCer1824 | 13.81 | 12.86 | 15.78 |
| R26 | DHCer1824.d -> DHCer1824 | 6.03 | 4.14 | 6.81 |
| R27 | DHCer1824 -> Cer1824.s | 19.84 | 18.39 | 21.34 |
| R28 | Spho18 + C24FA -> Cer1824.s | 25.18 | 21.79 | 30.93 |
| R29 | Cer1824.s -> Cer1824 | 45.02 | 41.77 | 50.66 |
| R30 | Cer1824.d -> Cer1824 | 55.95 | 50.45 | 62.08 |
| R31 | Spho18.s -> Spho18 | 15.14 | 10.55 | 20.87 |
| R32 | Spho18.d -> Spho18 | 10.05 | 6.64 | 13.44 |
| R33 | Cer1824 + Chol -> SM42_1.s | 100.97 | 93.34 | 109.45 |
| R34 | SM42_1.s -> SM42_1 | 100.97 | 93.34 | 109.45 |
| R35 | SM42_1.d -> SM42_1 | 14.17 | 7.37 | 20.64 |
| R36 | SM42_1 -> Spho18.s + C24FA.r + Chol.snk | 15.14 | 10.55 | 20.87 |
| R37 | SM42_1 -> SM42_1.m | 100.00 | 100.00 | 100.00 |
