## Supplementary material for "Modeling compound lipid homeostasis using stable isotope tracing": MFA results: A549 SL 20% FBS

### A549 20% FBS

SSR: 894.6 [740.6 899.2]

| Rxn# | Atom Transition | Flux (nanomoles) | Lower Bound | Upper Bound |
|---|---|---|---|---|
| R1 | Ac.l -> Ac.f | 44.43 | 41.84 | 47.20 |
| R2 | Ac.d -> Ac.f | 114.61 | 88.56 | 153.33 |
| R3 | Ac.f -> Ac | 159.03 | 134.11 | 196.17 |
| R4 | Ac -> AcER.l | 100.80 | 87.32 | 118.52 |
| R5 | AcER.l -> AcER.f | 100.80 | 87.32 | 118.52 |
| R6 | AcER.d -> AcER.f | 47.88 | 30.94 | 60.31 |
| R7 | AcER.f -> AcER | 148.68 | 142.22 | 158.39 |
| R8 | Ac + Ac + Ac + Ac + Ac + Ac + Ac + Ac -> Palm.s | 7.28 | 5.58 | 10.19 |
| R9 | Palm.s -> Palm | 7.28 | 5.58 | 10.19 |
| R10 | Palm.d -> Palm | 70.93 | 68.84 | 72.81 |
| R11 | Palm + AcER + AcER + AcER + AcER -> C24FA.s | 37.17 | 35.55 | 39.60 |
| R12 | C24FA.s -> C24FA | 37.17 | 35.55 | 39.60 |
| R13 | C24FA.d -> C24FA | 9.45 | 6.94 | 11.37 |
| R14 | C24FA.r -> C24FA | 1.16 | 0.78 | 1.58 |
| R15 | SerM1.l -> Ser.f | 7.89 | 6.92 | 8.85 |
| R16 | SerM2.l -> Ser.f | 4.00 | 1.89 | 6.05 |
| R17 | SerM3.l -> Ser.f | 104.44 | 102.14 | 106.83 |
| R18 | Ser.d -> Ser.f | 33.67 | 31.85 | 35.48 |
| R19 | Ser.f -> Ser | 150.00 | 150.00 | 150.00 |
| R20 | Ser -> Ser.snk | 108.96 | 107.91 | 109.99 |
| R21 | Palm + Ser -> Spha18.s + CO2 | 41.04 | 40.01 | 42.09 |
| R22 | Spha18.s -> Spha18 | 41.04 | 40.01 | 42.09 |
| R23 | Spha18.d -> Spha18 | 0.00 | 0.00 | 1.10 |
| R24 | Spha18 + C24FA -> DHCer1824.s | 41.04 | 40.01 | 42.42 |
| R25 | DHCer1824.s -> DHCer1824 | 41.04 | 40.01 | 42.42 |
| R26 | DHCer1824.d -> DHCer1824 | 7.77 | 6.54 | 8.59 |
| R27 | DHCer1824 -> Cer1824.s | 48.81 | 47.73 | 49.89 |
| R28 | Spho18 + C24FA -> Cer1824.s | 6.74 | 5.87 | 7.63 |
| R29 | Cer1824.s -> Cer1824 | 55.55 | 54.37 | 56.73 |
| R30 | Cer1824.d -> Cer1824 | 0.00 | 0.00 | 0.19 |
| R31 | Spho18.s -> Spho18 | 1.16 | 0.78 | 1.58 |
| R32 | Spho18.d -> Spho18 | 5.58 | 4.85 | 6.31 |
| R33 | Cer1824 + Chol -> SM42_1.s | 55.55 | 54.37 | 56.73 |
| R34 | SM42_1.s -> SM42_1 | 55.55 | 54.37 | 56.73 |
| R35 | SM42_1.d -> SM42_1 | 45.61 | 44.48 | 46.75 |
| R36 | SM42_1 -> Spho18.s + C24FA.r + Chol.snk | 1.16 | 0.78 | 1.58 |
| R37 | SM42_1 -> SM42_1.m | 100.00 | 100.00 | 100.00 |
