## Supplementary material for "Modeling compound lipid homeostasis using stable isotope tracing": MFA results: A549 20% FBS +veh

| | **A549 20% FBS +Vehicle** | | 99% Confidence Intervals | |
|---|---|---|---|---|
| Rxn# | Atom Transition | Flux (nanomoles) | Lower Bound | Upper Bound |
| R1 | Ac.l -> Ac.f | 104.88 | 100.16 | 109.70 |
| R2 | Ac.d -> Ac.f | 437.24 | 418.06 | 456.94 |
| R3 | Ac.f -> Ac | 542.12 | 520.37 | 564.27 |
| R4 | Ac -> AcER.l | 86.31 | 79.22 | 94.31 |
| R5 | AcER.l -> AcER.f | 86.31 | 79.22 | 94.31 |
| R6 | AcER.d -> AcER.f | 5.23 | 0.00 | 10.57 |
| R7 | AcER.f -> AcER | 91.54 | 84.01 | 100.40 |
| R8 | glyc.l -> glyc.f | 188.26 | 186.66 | 189.85 |
| R9 | glyc.d -> glyc.f | 102.10 | 98.11 | 105.34 |
| R10 | glyc.f -> glyc | 290.36 | 286.25 | 293.98 |
| R11 | Ac + Ac + Ac + Ac + Ac + Ac + Ac + Ac -> Palm.s | 56.11 | 53.82 | 58.43 |
| R12 | Palm.s -> Palm | 56.11 | 53.82 | 58.43 |
| R13 | Palm.d -> Palm | 292.82 | 281.19 | 302.56 |
| R14 | Palm.r -> Palm | 0.00 | 0.00 | 6.05 |
| R15 | Palm + AcER -> Stea.s | 48.79 | 41.80 | 57.32 |
| R16 | Stea.s -> Stea | 48.79 | 41.80 | 57.32 |
| R17 | Stea.d -> Stea | 36.76 | 30.24 | 45.55 |
| R18 | Stea -> Olea.s | 9.10 | 0.00 | 24.88 |
| R19 | Olea.s -> Olea | 9.10 | 0.00 | 24.88 |
| R20 | Olea.d -> Olea | 266.73 | 251.20 | 278.25 |
| R21 | Stea + AcER + AcER + AcER -> C24_0FA.s | 6.80 | 6.64 | 6.97 |
| R22 | C24_0FA.s -> C24_0FA | 6.80 | 6.64 | 6.97 |
| R23 | C24_0FA.d -> C24_0FA | 0.42 | 0.00 | 0.64 |
| R24 | C24_0FA.r -> C24_0FA | 1.94 | 0.00 | 2.85 |
| R25 | SerM1.l -> Ser.f | 14.39 | 12.89 | 15.88 |
| R26 | SerM2.l -> Ser.f | 3.86 | 0.58 | 7.09 |
| R27 | SerM3.l -> Ser.f | 270.70 | 266.70 | 275.00 |
| R28 | Ser.d -> Ser.f | 97.04 | 93.32 | 100.37 |
| R29 | Ser.f -> Ser | 386.00 | 386.00 | 386.00 |
| R30 | Ser -> Ser.snk | 325.56 | 324.85 | 326.42 |
| R31 | Palm + Ser -> Spha18.s + CO2 | 33.76 | 33.12 | 34.40 |
| R32 | Spha18.s -> Spha18 | 33.76 | 33.12 | 34.40 |
| R33 | Spha18.d -> Spha18 | 0.00 | 0.00 | 0.27 |
| R34 | Spha18 + C24_0FA -> DHCer1824_0.s | 8.13 | 7.98 | 8.51 |
| R35 | DHCer1824_0.s -> DHCer1824_0 | 8.13 | 7.98 | 8.51 |
| R36 | DHCer1824_0.d -> DHCer1824_0 | 1.24 | 1.09 | 1.40 |
| R37 | DHCer1824_0 -> Cer1824_0.s | 9.23 | 9.05 | 9.66 |
| R38 | Spho18 + C24_0FA -> Cer1824_0.s | 1.03 | 0.87 | 1.22 |

| | | | | |
|---|---|---|---|---|
| R39 | Cer1824_0.s -> Cer1824_0 | 10.26 | 10.12 | 10.77 |
| R40 | Cer1824_0.d -> Cer1824_0 | 0.00 | 0.00 | 0.02 |
| R41 | Spho18.s -> Spho18 | 1.94 | 1.17 | 3.92 |
| R42 | Spho18.d -> Spho18 | 8.97 | 6.62 | 11.25 |
| R43 | Cer1824_0 + Chol -> SM42_1.s | 5.95 | 5.86 | 6.44 |
| R44 | SM42_1.s -> SM42_1 | 5.95 | 5.86 | 6.44 |
| R45 | SM42_1.d -> SM42_1 | 5.41 | 5.32 | 5.85 |
| R46 | SM42_1 -> Spho18.s + C24_0FA.r + Chol.snk | 1.94 | 0.00 | 2.85 |
| R47 | SM42_1 -> SM42_1.m | 9.41 | 9.41 | 9.41 |
| R48 | glyc + Palm + Palm -> DG32.s | 21.98 | 21.68 | 22.24 |
| R49 | DG32.s -> DG32 | 21.98 | 21.68 | 22.24 |
| R50 | DG32.d -> DG32 | 3.94 | 0.00 | 4.24 |
| R51 | DG32 + Chol -> PC32.s | 25.91 | 25.61 | 27.92 |
| R52 | PC32.s -> PC32 | 25.91 | 25.61 | 27.92 |
| R53 | PC32.d -> PC32 | 2.01 | 0.00 | 2.30 |
| R54 | PC32 -> PC32.m | 27.92 | 27.92 | 27.92 |
| R55 | glyc + Palm + Olea -> DG34.s | 198.74 | 195.42 | 201.76 |
| R56 | DG34.s -> DG34 | 198.74 | 195.42 | 201.76 |
| R57 | DG34.d -> DG34 | 0.00 | 0.00 | 3.13 |
| R58 | DG34 + Chol -> PC34.s | 192.30 | 189.31 | 195.45 |
| R59 | PC34.s -> PC34 | 192.30 | 189.31 | 195.45 |
| R60 | PC34.d -> PC34 | 141.88 | 138.72 | 144.86 |
| R61 | PC34 -> PC34.m | 334.17 | 334.17 | 334.17 |
| R62 | DG34 + Ser -> PS34_1.s | 5.45 | 5.35 | 5.56 |
| R63 | PS34_1.s -> PS34_1 | 5.45 | 5.35 | 5.56 |
| R64 | PS34_1.d -> PS34_1 | 0.36 | 0.26 | 0.46 |
| R65 | PS34_1 -> PS34_1.m | 4.42 | 4.42 | 4.42 |
| R66 | PS34_1 -> PE34.s + CO2 | 1.39 | 1.32 | 1.45 |
| R67 | DG34 + EtA -> PE34.s | 0.99 | 0.92 | 1.06 |
| R68 | PE34.s -> PE34 | 2.38 | 2.34 | 2.42 |
| R69 | PE34.d -> PE34 | 1.54 | 1.50 | 1.58 |
| R70 | PE34 -> PE34.m | 3.92 | 3.92 | 3.92 |
| R71 | glyc + Stea + Olea -> DG36.s | 69.64 | 68.75 | 70.47 |
| R72 | DG36.s -> DG36 | 69.64 | 68.75 | 70.47 |
| R73 | DG36.d -> DG36 | 0.00 | 0.00 | 0.43 |
| R74 | DG36 + Chol -> PC36.s | 47.23 | 46.38 | 48.02 |
| R75 | PC36.s -> PC36 | 47.23 | 46.38 | 48.02 |
| R76 | PC36.d -> PC36 | 41.18 | 40.39 | 42.03 |
| R77 | PC36 -> PC36.m | 88.41 | 88.41 | 88.41 |
| R78 | DG36 + Ser -> PS36_1.s | 21.23 | 20.93 | 21.29 |
| R79 | PS36_1.s -> PS36_1 | 21.23 | 20.93 | 21.29 |

| | | | | |
|---|---|---|---|---|
| R80 | PS36_1.d -> PS36_1 | 0.00 | 0.00 | 0.33 |
| R81 | PS36_1 -> PS36_1.m | 17.67 | 17.67 | 17.67 |
| R82 | PS36_1 -> PE36.s + CO2 | 3.56 | 3.50 | 3.62 |
| R83 | DG36 + EtA -> PE36.s | 1.19 | 1.09 | 1.28 |
| R84 | PE36.s -> PE36 | 4.75 | 4.67 | 4.83 |
| R85 | PE36.d -> PE36 | 4.02 | 3.94 | 4.10 |
| R86 | PE36 -> PE36.m | 8.77 | 8.77 | 8.77 |
| R87 | Spha18 + Palm -> DHCer1816.s | 17.13 | 16.56 | 17.68 |
| R88 | DHCer1816.s -> DHCer1816 | 17.13 | 16.56 | 17.68 |
| R89 | DHCer1816.d -> DHCer1816 | 11.75 | 10.96 | 12.91 |
| R90 | DHCer1816 -> Cer1816.s | 20.23 | 19.22 | 21.38 |
| R91 | Spho18 + Palm -> Cer1816.s | 6.55 | 4.02 | 10.02 |
| R92 | Cer1816.s -> Cer1816 | 26.78 | 24.38 | 30.86 |
| R93 | Cer1816.d -> Cer1816 | 4.75 | 2.55 | 7.08 |
| R94 | Cer1816 + Chol -> SM34_1.s | 30.54 | 28.97 | 33.29 |
| R95 | SM34_1.s -> SM34_1 | 30.54 | 28.97 | 33.29 |
| R96 | SM34_1.d -> SM34_1 | 27.67 | 26.03 | 31.40 |
| R97 | SM34_1 -> Spho18.s + Palm.r + Chol.snk | 0.00 | 0.00 | 6.05 |
| R98 | SM34_1 -> SM34_1.m | 58.21 | 58.21 | 58.21 |
| R99 | Olea + AcER + AcER + AcER -> C24_1FA.s | 7.45 | 7.21 | 7.71 |
| R100 | C24_1FA.s -> C24_1FA | 7.45 | 7.21 | 7.71 |
| R101 | C24_1FA.d -> C24_1FA | 4.38 | 3.77 | 4.81 |
| R102 | C24_1FA.r -> C24_1FA | 0.00 | 0.00 | 1.02 |
| R103 | Spha18 + C24_1FA -> DHCer1824_1.s | 8.50 | 8.30 | 8.86 |
| R104 | DHCer1824_1.s -> DHCer1824_1 | 8.50 | 8.30 | 8.86 |
| R105 | DHCer1824_1.d -> DHCer1824_1 | 0.20 | 0.03 | 0.37 |
| R106 | DHCer1824_1 -> Cer1824_1.s | 8.70 | 8.46 | 9.08 |
| R107 | Spho18 + C24_1FA -> Cer1824_1.s | 3.33 | 2.92 | 3.82 |
| R108 | Cer1824_1.s -> Cer1824_1 | 12.03 | 11.71 | 12.69 |
| R109 | Cer1824_1.d -> Cer1824_1 | 0.45 | 0.20 | 0.69 |
| R110 | Cer1824_1 + Chol -> SM42_2.s | 12.48 | 12.24 | 13.11 |
| R111 | SM42_2.s -> SM42_2 | 12.48 | 12.24 | 13.11 |
| R112 | SM42_2.d -> SM42_2 | 9.79 | 9.54 | 10.29 |
| R113 | SM42_2 -> Spho18.s + C24_1FA.r + Chol.snk | 0.00 | 0.00 | 1.02 |
| R114 | SM42_2 -> SM42_2.m | 22.27 | 22.27 | 22.27 |
| R115 | EtA.d -> EtA | 2.18 | 2.05 | 2.31 |
| R116 | DHCer1824_0 + Chol -> DHSM42_0.s | 0.14 | 0.14 | 0.15 |
| R117 | DHSM42_0.s -> DHSM42_0 | 0.14 | 0.14 | 0.15 |
| R118 | DHSM42_0.d -> DHSM42_0 | 0.10 | 0.09 | 0.10 |
| R119 | DHSM42_0 -> DHSM42_0.m | 0.24 | 0.24 | 0.24 |
| R120 | DHCer1816 + Chol -> DHSM34_0.s | 8.65 | 8.42 | 8.89 |

| | | | | |
|---|---|---|---|---|
| R121 | DHSM34_0.s -> DHSM34_0 | 8.65 | 8.42 | 8.89 |
| R122 | DHSM34_0.d -> DHSM34_0 | 0.93 | 0.70 | 1.16 |
| R123 | DHSM34_0 -> DHSM34_0.m | 9.58 | 9.58 | 9.58 |
| R124 | Glc.l -> Glc | 15.91 | 15.46 | 16.33 |
| R125 | Glc.d -> Glc | 0.00 | 0.00 | 1.89 |
| R126 | Glc -> Gal.s | 3.64 | 3.17 | 4.99 |
| R127 | Gal.s -> Gal | 3.64 | 3.17 | 4.99 |
| R128 | Gal.d -> Gal | 1.67 | 0.32 | 2.14 |
| R129 | PEP.l -> PEP | 3.32 | 3.15 | 3.49 |
| R130 | PEP.d -> PEP | 0.30 | 0.00 | 0.61 |
| R131 | Glc + Ac -> GlcNac.s | 6.96 | 6.38 | 7.86 |
| R132 | GlcNac.s -> GlcNac | 6.96 | 6.38 | 7.86 |
| R133 | GlcNac.d -> GlcNac | 2.09 | 1.22 | 2.69 |
| R134 | GlcNac + PEP -> NeuAc.s | 3.62 | 3.36 | 3.89 |
| R135 | NeuAc.s -> NeuAc | 3.62 | 3.36 | 3.89 |
| R136 | NeuAc.d -> NeuAc | 1.69 | 1.44 | 1.93 |
| R137 | Cer1816 + Glc -> HexCer34_1.s | 1.00 | 0.96 | 1.04 |
| R138 | HexCer34_1.s -> HexCer34_1 | 1.00 | 0.96 | 1.04 |
| R139 | HexCer34_1.d -> HexCer34_1 | 0.00 | 0.00 | 0.03 |
| R140 | HexCer34_1 + Gal -> LacCer.s | 1.00 | 0.96 | 1.04 |
| R141 | LacCer.s -> LacCer | 1.00 | 0.96 | 1.04 |
| R142 | LacCer.d -> LacCer | 0.00 | 0.00 | 0.05 |
| R143 | LacCer + NeuAc -> GM3_34_1.s | 1.00 | 0.96 | 1.04 |
| R144 | GM3_34_1.s -> GM3_34_1 | 1.00 | 0.96 | 1.04 |
| R145 | GM3_34_1.d -> GM3_34_1 | 0.08 | 0.05 | 0.11 |
| R146 | GM3_34_1 + GlcNac -> GM2_34_1.s | 1.08 | 1.04 | 1.12 |
| R147 | GM2_34_1.s -> GM2_34_1 | 1.08 | 1.04 | 1.12 |
| R148 | GM2_34_1.d -> GM2_34_1 | 1.67 | 1.63 | 1.71 |
| R149 | GM2_34_1 -> GM2_34_1.m | 2.75 | 2.75 | 2.75 |
| R150 | Cer1824_0 + Glc -> HexCer42_1.s | 4.31 | 4.21 | 4.41 |
| R151 | HexCer42_1.s -> HexCer42_1 | 4.31 | 4.21 | 4.41 |
| R152 | HexCer42_1.d -> HexCer42_1 | 0.00 | 0.00 | 0.12 |
| R153 | HexCer42_1 + Gal -> LacCer42_1.s | 4.31 | 4.21 | 4.44 |
| R154 | LacCer42_1.s -> LacCer42_1 | 4.31 | 4.21 | 4.44 |
| R155 | LacCer42_1.d -> LacCer42_1 | 0.00 | 0.00 | 0.09 |
| R156 | LacCer42_1 + NeuAc -> GM3_42_1.s | 4.31 | 4.21 | 4.44 |
| R157 | GM3_42_1.s -> GM3_42_1 | 4.31 | 4.21 | 4.44 |
| R158 | GM3_42_1.d -> GM3_42_1 | 0.04 | 0.00 | 0.15 |
| R159 | GM3_42_1 + GlcNac -> GM2_42_1.s | 4.35 | 4.23 | 4.49 |
| R160 | GM2_42_1.s -> GM2_42_1 | 4.35 | 4.23 | 4.49 |
| R161 | GM2_42_1.d -> GM2_42_1 | 6.40 | 6.26 | 6.52 |

| | | | | |
|---|---|---|---|---|
| R162 | GM2_42_1 -> GM2_42_1.m | 10.75 | 10.75 | 10.75 |
