## Supplementary material for "Modeling compound lipid homeostasis using stable isotope tracing": MFA results: A549 20% FBS + FuB1

| | **A549 20% FBS +2uM Fub1** | | 99% Confidence Intervals | |
|---|---|---|---|---|
| Rxn# | Atom Transition | Flux (nanomoles) | Lower Bound | Upper Bound |
| R1 | Ac.l -> Ac.f | 117.06 | 114.50 | 119.65 |
| R2 | Ac.d -> Ac.f | 452.35 | 441.80 | 462.97 |
| R3 | Ac.f -> Ac | 569.41 | 558.27 | 580.56 |
| R4 | Ac -> AcER.l | 116.45 | 113.63 | 119.39 |
| R5 | AcER.l -> AcER.f | 116.45 | 113.63 | 119.39 |
| R6 | AcER.d -> AcER.f | 3.86 | 0.11 | 7.64 |
| R7 | AcER.f -> AcER | 120.31 | 117.53 | 123.25 |
| R8 | glyc.l -> glyc.f | 191.61 | 190.36 | 192.86 |
| R9 | glyc.d -> glyc.f | 85.56 | 83.00 | 88.14 |
| R10 | glyc.f -> glyc | 277.17 | 274.29 | 280.07 |
| R11 | Ac + Ac + Ac + Ac + Ac + Ac + Ac + Ac -> Palm.s | 55.63 | 54.35 | 56.91 |
| R12 | Palm.s -> Palm | 55.63 | 54.35 | 56.91 |
| R13 | Palm.d -> Palm | 282.35 | 271.81 | 287.26 |
| R14 | Palm.r -> Palm | 0.00 | 0.00 | 11.55 |
| R15 | Palm + AcER -> Stea.s | 45.59 | 43.51 | 47.75 |
| R16 | Stea.s -> Stea | 45.59 | 43.51 | 47.75 |
| R17 | Stea.d -> Stea | 31.90 | 30.13 | 33.64 |
| R18 | Stea -> Olea.s | 0.00 | 0.00 | 1.25 |
| R19 | Olea.s -> Olea | 0.00 | 0.00 | 1.25 |
| R20 | Olea.d -> Olea | 263.88 | 261.13 | 266.67 |
| R21 | Stea + AcER + AcER + AcER -> C24_0FA.s | 13.13 | 12.95 | 13.33 |
| R22 | C24_0FA.s -> C24_0FA | 13.13 | 12.95 | 13.33 |
| R23 | C24_0FA.d -> C24_0FA | 1.49 | 1.07 | 2.60 |
| R24 | C24_0FA.r -> C24_0FA | 2.30 | 0.00 | 3.22 |
| R25 | SerM1.l -> Ser.f | 16.46 | 15.05 | 17.96 |
| R26 | SerM2.l -> Ser.f | 2.38 | 0.00 | 5.97 |
| R27 | SerM3.l -> Ser.f | 330.68 | 311.18 | 351.80 |
| R28 | Ser.d -> Ser.f | 36.48 | 14.52 | 56.76 |
| R29 | Ser.f -> Ser | 386.00 | 386.00 | 386.00 |
| R30 | Ser -> Ser.snk | 330.81 | 327.39 | 334.08 |
| R31 | Palm + Ser -> Spha18.s + CO2 | 39.58 | 37.22 | 42.05 |
| R32 | Spha18.s -> Spha18 | 39.58 | 37.22 | 42.05 |
| R33 | Spha18.d -> Spha18 | 2.44 | 0.00 | 4.78 |
| R34 | Spha18 + C24_0FA -> DHCer1824_0.s | 15.75 | 15.58 | 16.28 |
| R35 | DHCer1824_0.s -> DHCer1824_0 | 15.75 | 15.58 | 16.28 |
| R36 | DHCer1824_0.d -> DHCer1824_0 | 2.04 | 1.89 | 2.21 |
| R37 | DHCer1824_0 -> Cer1824_0.s | 17.22 | 17.02 | 17.83 |
| R38 | Spho18 + C24_0FA -> Cer1824_0.s | 1.18 | 0.99 | 1.38 |

| | | | | |
|---|---|---|---|---|
| R39 | Cer1824_0.s -> Cer1824_0 | 18.40 | 18.24 | 19.05 |
| R40 | Cer1824_0.d -> Cer1824_0 | 0.00 | 0.00 | 0.01 |
| R41 | Spho18.s -> Spho18 | 2.30 | 1.41 | 6.02 |
| R42 | Spho18.d -> Spho18 | 5.24 | 0.00 | 7.29 |
| R43 | Cer1824_0 + Chol -> SM42_1.s | 12.95 | 12.83 | 13.59 |
| R44 | SM42_1.s -> SM42_1 | 12.95 | 12.83 | 13.59 |
| R45 | SM42_1.d -> SM42_1 | 5.92 | 5.80 | 6.23 |
| R46 | SM42_1 -> Spho18.s + C24_0FA.r + Chol.snk | 2.30 | 0.00 | 3.22 |
| R47 | SM42_1 -> SM42_1.m | 16.57 | 16.57 | 16.57 |
| R48 | glyc + Palm + Palm -> DG32.s | 25.06 | 24.86 | 25.26 |
| R49 | DG32.s -> DG32 | 25.06 | 24.86 | 25.26 |
| R50 | DG32.d -> DG32 | 7.36 | 7.14 | 7.56 |
| R51 | DG32 + Chol -> PC32.s | 32.42 | 32.27 | 32.42 |
| R52 | PC32.s -> PC32 | 32.42 | 32.27 | 32.42 |
| R53 | PC32.d -> PC32 | 0.00 | 0.00 | 0.15 |
| R54 | PC32 -> PC32.m | 32.42 | 32.42 | 32.42 |
| R55 | glyc + Palm + Olea -> DG34.s | 187.75 | 185.49 | 190.03 |
| R56 | DG34.s -> DG34 | 187.75 | 185.49 | 190.03 |
| R57 | DG34.d -> DG34 | 3.96 | 1.48 | 6.45 |
| R58 | DG34 + Chol -> PC34.s | 187.99 | 185.45 | 190.55 |
| R59 | PC34.s -> PC34 | 187.99 | 185.45 | 190.55 |
| R60 | PC34.d -> PC34 | 145.83 | 143.27 | 148.36 |
| R61 | PC34 -> PC34.m | 333.81 | 333.81 | 333.81 |
| R62 | DG34 + Ser -> PS34_1.s | 2.60 | 2.45 | 2.77 |
| R63 | PS34_1.s -> PS34_1 | 2.60 | 2.45 | 2.77 |
| R64 | PS34_1.d -> PS34_1 | 0.66 | 0.50 | 0.82 |
| R65 | PS34_1 -> PS34_1.m | 1.79 | 1.79 | 1.79 |
| R66 | PS34_1 -> PE34.s + CO2 | 1.48 | 1.43 | 1.53 |
| R67 | DG34 + EtA -> PE34.s | 1.12 | 1.03 | 1.21 |
| R68 | PE34.s -> PE34 | 2.60 | 2.51 | 2.68 |
| R69 | PE34.d -> PE34 | 1.29 | 1.21 | 1.38 |
| R70 | PE34 -> PE34.m | 3.89 | 3.89 | 3.89 |
| R71 | glyc + Stea + Olea -> DG36.s | 64.36 | 63.44 | 65.29 |
| R72 | DG36.s -> DG36 | 64.36 | 63.44 | 65.29 |
| R73 | DG36.d -> DG36 | 0.00 | 0.00 | 0.15 |
| R74 | DG36 + Chol -> PC36.s | 49.92 | 49.22 | 50.62 |
| R75 | PC36.s -> PC36 | 49.92 | 49.22 | 50.62 |
| R76 | PC36.d -> PC36 | 47.20 | 46.50 | 47.90 |
| R77 | PC36 -> PC36.m | 97.12 | 97.12 | 97.12 |
| R78 | DG36 + Ser -> PS36_1.s | 13.01 | 12.23 | 13.82 |
| R79 | PS36_1.s -> PS36_1 | 13.01 | 12.23 | 13.82 |

| | | | | |
|---|---|---|---|---|
| R80 | PS36_1.d -> PS36_1 | 2.39 | 1.57 | 3.18 |
| R81 | PS36_1 -> PS36_1.m | 10.74 | 10.74 | 10.74 |
| R82 | PS36_1 -> PE36.s + CO2 | 4.66 | 4.59 | 4.73 |
| R83 | DG36 + EtA -> PE36.s | 1.43 | 1.18 | 1.68 |
| R84 | PE36.s -> PE36 | 6.09 | 5.83 | 6.34 |
| R85 | PE36.d -> PE36 | 4.36 | 4.10 | 4.61 |
| R86 | PE36 -> PE36.m | 10.44 | 10.44 | 10.44 |
| R87 | Spha18 + Palm -> DHCer1816.s | 12.19 | 11.51 | 12.83 |
| R88 | DHCer1816.s -> DHCer1816 | 12.19 | 11.51 | 12.83 |
| R89 | DHCer1816.d -> DHCer1816 | 13.45 | 12.65 | 14.05 |
| R90 | DHCer1816 -> Cer1816.s | 15.19 | 13.77 | 16.55 |
| R91 | Spho18 + Palm -> Cer1816.s | 2.75 | 0.00 | 7.07 |
| R92 | Cer1816.s -> Cer1816 | 17.95 | 16.13 | 21.76 |
| R93 | Cer1816.d -> Cer1816 | 5.01 | 3.05 | 7.53 |
| R94 | Cer1816 + Chol -> SM34_1.s | 22.55 | 21.48 | 24.83 |
| R95 | SM34_1.s -> SM34_1 | 22.55 | 21.48 | 24.83 |
| R96 | SM34_1.d -> SM34_1 | 22.23 | 21.15 | 28.56 |
| R97 | SM34_1 -> Spho18.s + Palm.r + Chol.snk | 0.00 | 0.00 | 11.55 |
| R98 | SM34_1 -> SM34_1.m | 44.77 | 44.77 | 44.77 |
| R99 | Olea + AcER + AcER + AcER -> C24_1FA.s | 11.77 | 11.57 | 11.99 |
| R100 | C24_1FA.s -> C24_1FA | 11.77 | 11.57 | 11.99 |
| R101 | C24_1FA.d -> C24_1FA | 5.92 | 5.39 | 6.14 |
| R102 | C24_1FA.r -> C24_1FA | 0.00 | 0.00 | 1.15 |
| R103 | Spha18 + C24_1FA -> DHCer1824_1.s | 14.08 | 13.88 | 14.62 |
| R104 | DHCer1824_1.s -> DHCer1824_1 | 14.08 | 13.88 | 14.62 |
| R105 | DHCer1824_1.d -> DHCer1824_1 | 0.00 | 0.00 | 0.05 |
| R106 | DHCer1824_1 -> Cer1824_1.s | 14.08 | 13.88 | 14.62 |
| R107 | Spho18 + C24_1FA -> Cer1824_1.s | 3.62 | 3.35 | 3.93 |
| R108 | Cer1824_1.s -> Cer1824_1 | 17.70 | 17.49 | 18.40 |
| R109 | Cer1824_1.d -> Cer1824_1 | 0.56 | 0.40 | 0.72 |
| R110 | Cer1824_1 + Chol -> SM42_2.s | 18.26 | 18.08 | 18.96 |
| R111 | SM42_2.s -> SM42_2 | 18.26 | 18.08 | 18.96 |
| R112 | SM42_2.d -> SM42_2 | 12.90 | 12.72 | 13.41 |
| R113 | SM42_2 -> Spho18.s + C24_1FA.r + Chol.snk | 0.00 | 0.00 | 1.15 |
| R114 | SM42_2 -> SM42_2.m | 31.16 | 31.16 | 31.16 |
| R115 | EtA.d -> EtA | 2.55 | 2.22 | 2.87 |
| R116 | DHCer1824_0 + Chol -> DHSM42_0.s | 0.57 | 0.56 | 0.58 |
| R117 | DHSM42_0.s -> DHSM42_0 | 0.57 | 0.56 | 0.58 |
| R118 | DHSM42_0.d -> DHSM42_0 | 0.17 | 0.16 | 0.18 |
| R119 | DHSM42_0 -> DHSM42_0.m | 0.74 | 0.74 | 0.74 |
| R120 | DHCer1816 + Chol -> DHSM34_0.s | 10.44 | 10.27 | 10.44 |

| | | | | |
|---|---|---|---|---|
| R121 | DHSM34_0.s -> DHSM34_0 | 10.44 | 10.27 | 10.44 |
| R122 | DHSM34_0.d -> DHSM34_0 | 0.00 | 0.00 | 0.18 |
| R123 | DHSM34_0 -> DHSM34_0.m | 10.44 | 10.44 | 10.44 |
| R124 | Glc.l -> Glc | 17.37 | 16.96 | 17.79 |
| R125 | Glc.d -> Glc | 0.00 | 0.00 | 0.72 |
| R126 | Glc -> Gal.s | 3.57 | 3.02 | 5.00 |
| R127 | Gal.s -> Gal | 3.57 | 3.02 | 5.00 |
| R128 | Gal.d -> Gal | 2.29 | 0.82 | 2.82 |
| R129 | PEP.l -> PEP | 3.98 | 3.81 | 4.15 |
| R130 | PEP.d -> PEP | 0.25 | 0.00 | 0.57 |
| R131 | Glc + Ac -> GlcNac.s | 7.95 | 6.77 | 8.61 |
| R132 | GlcNac.s -> GlcNac | 7.95 | 6.77 | 8.61 |
| R133 | GlcNac.d -> GlcNac | 2.33 | 1.65 | 3.61 |
| R134 | GlcNac + PEP -> NeuAc.s | 4.23 | 3.97 | 4.52 |
| R135 | NeuAc.s -> NeuAc | 4.23 | 3.97 | 4.52 |
| R136 | NeuAc.d -> NeuAc | 1.63 | 1.33 | 1.88 |
| R137 | Cer1816 + Glc -> HexCer34_1.s | 0.41 | 0.35 | 0.44 |
| R138 | HexCer34_1.s -> HexCer34_1 | 0.41 | 0.35 | 0.44 |
| R139 | HexCer34_1.d -> HexCer34_1 | 0.00 | 0.00 | 0.06 |
| R140 | HexCer34_1 + Gal -> LacCer.s | 0.41 | 0.37 | 0.45 |
| R141 | LacCer.s -> LacCer | 0.41 | 0.37 | 0.45 |
| R142 | LacCer.d -> LacCer | 0.01 | 0.00 | 0.04 |
| R143 | LacCer + NeuAc -> GM3_34_1.s | 0.41 | 0.38 | 0.45 |
| R144 | GM3_34_1.s -> GM3_34_1 | 0.41 | 0.38 | 0.45 |
| R145 | GM3_34_1.d -> GM3_34_1 | 0.09 | 0.06 | 0.12 |
| R146 | GM3_34_1 + GlcNac -> GM2_34_1.s | 0.50 | 0.47 | 0.54 |
| R147 | GM2_34_1.s -> GM2_34_1 | 0.50 | 0.47 | 0.54 |
| R148 | GM2_34_1.d -> GM2_34_1 | 1.82 | 1.78 | 1.85 |
| R149 | GM2_34_1 -> GM2_34_1.m | 2.32 | 2.32 | 2.32 |
| R150 | Cer1824_0 + Glc -> HexCer42_1.s | 5.45 | 5.35 | 5.55 |
| R151 | HexCer42_1.s -> HexCer42_1 | 5.45 | 5.35 | 5.55 |
| R152 | HexCer42_1.d -> HexCer42_1 | 0.00 | 0.00 | 0.03 |
| R153 | HexCer42_1 + Gal -> LacCer42_1.s | 5.45 | 5.35 | 5.55 |
| R154 | LacCer42_1.s -> LacCer42_1 | 5.45 | 5.35 | 5.55 |
| R155 | LacCer42_1.d -> LacCer42_1 | 0.00 | 0.00 | 0.02 |
| R156 | LacCer42_1 + NeuAc -> GM3_42_1.s | 5.45 | 5.35 | 5.55 |
| R157 | GM3_42_1.s -> GM3_42_1 | 5.45 | 5.35 | 5.55 |
| R158 | GM3_42_1.d -> GM3_42_1 | 0.09 | 0.00 | 0.20 |
| R159 | GM3_42_1 + GlcNac -> GM2_42_1.s | 5.54 | 5.42 | 5.66 |
| R160 | GM2_42_1.s -> GM2_42_1 | 5.54 | 5.42 | 5.66 |
| R161 | GM2_42_1.d -> GM2_42_1 | 7.23 | 7.11 | 7.35 |

| | | | | |
|---|---|---|---|---|
| R162 | GM2_42_1 -> GM2_42_1.m | 12.77 | 12.77 | 12.77 |
