## Supplementary material for "Modeling compound lipid homeostasis using stable isotope tracing": Transition Tables for LCMS

Table S1. Orbitrap high-resolution QE Mass spectrometry lipid analysis

| Lipid Species | Precursor Ion | Product Ion |
|---|---|---|
| **Long chain sphingoid bases (SPB)** | | |
| SBP 18:0;O2 | 302.3054 | 284.2974 |
| SBP 18:0;O2 [D7] | 309.3493 | 291.3401 |
| SBP 18:1;O2 | 300.2897 | 282.2782 |
| SBP 18:1;O2[D7] | 307.3336 | 289.3222 |
| **Dihydroceramides** | | |
| Cer 18:0;O2/16:0 | 540.535 | 266.3 |
| Cer 18:0;O2/24:0 | 652.6602 | 266.3 |
| Cer 18:0;O2/24:1 | 650.6446 | 266.3 |
| **Ceramides** | | |
| Cer 18:1;O2 [D7] /15:0 | 531.5477 | 271.3 |
| Cer 18:1;O2 [D7] /16:1 | 543.5477 | 271.3 |
| Cer 18:1;O2/16:0 | 538.5193 | 264.3 |
| Cer 18:1;O2 [D7] /24:1 | 655.6729 | 271.3 |
| Cer 18:1;O2/24:0 | 650.6446 | 264.3 |
| Cer 18:1;O2/24:1 | 648.6289 | 264.3 |
| **Sphingomyelin (SM)** | | |
| SM 18:1;O2/18:1 [D9] | 738.6443 | 184.0734 |
| SM 18:0;O2/16:0 | 705.5905 | 184.0734 |
| SM 18:1;O2/16:1 | 703.5748 | 184.0734 |
| SM 18:1;O2/16:1 [D9] | 710.6157 | 193.1296 |
| SM 18:1;O2/24:1 [D9] | 822.7409 | 193.1296 |
| SM 18:0;O2/24:0 | 817.7157 | 184.0734 |
| SM 18:1;O2/24:0 | 815.7001 | 184.0734 |
| SM 18:1;O2/24:1 | 813.6844 | 184.0734 |
| **Gangliosides (GM2/3)** | | |
| NeuAcHex2Cer 18:1;O2/18:0 [D3] | 1184.7706 | 264.3 |
| NeuAcHex2Cer 18:1;O2/16:0 | 1153.7205 | 264.3 |
| NeuAcHex2Cer 18:1;O2/24:0 | 1265.8456 | 264.3 |
| HexNAcNeuAcHex2Cer 18:1;O2/16:0 [D9] | 1365.8628 | 264.3 |
| HexNAcNeuAcHex2Cer 18:1;O2/16:0 | 1356.7998 | 264.3 |

| | | |
|---|---|---|
| HexNAcNeuAcHex2Cer 18:1;O2/24:0 | 1265.8456 | 264.3 |
| **Globosides (GB3)** | | |
| Hex3Cer 18:1;O2/16:0 [D9] | 1033.7332 | 264.3 |
| Hex3Cer 18:1;O2/16:0 | 1024.6779 | 264.3 |
| Hex3Cer 18:1;O2/24:0 | 1136.8031 | 264.3 |
| **Diacylglycerols (DG)** | | |
| DG 15:0_18:1 [D7] | 605.5858 | 346.3 |
| DG 16:0_16:0 | 586.5417 | 313.3 |
| DG 16:0_18:1 | 612.5576 | 313.3 |
| DG 18:0_18:1 | 640.589 | 341.3 |
| **Triacylglycerols (TG)** | | |
| TG 15:0_18:1 [D7]_15:0 | 829.7999 | 570.5 |
| TG 16:0_18:1_18:0 | 878.8138 | 579.5 |
| TG 24:0_32:0 | 936.8954 | 549.4965 (FA24:0 loss) |
| TG 24:0_32:1<br>TG 24:1_32:0 | 934.8797 | 547.8372 (FA24:0 loss)<br>549.4869 (FA24:1 loss) |
| TG 24:1_32:1 | 932.8641 | 549.4965 (FA24:1 loss) |
| TG 24:0_34:2<br>TG 24:1_34:1 | 960.8954 | 575.5037 (FA24:0 loss)<br>577.5171 (FA24:1 loss) |
| TG 24:1_34:2 | 958.8812 | 575.5037 (FA24:1 loss) |
| TG 24:0_36:1 | 990.9423 | 605.5502 (FA24:0 loss) |
| TG 24:0_36:2<br>TG 24:1_36:1 | 988.9266 | 603.5336 (FA24:0 loss)<br>605.5502 (FA24:1 loss) |
| **Phosphatidylcholines (PC)** | | |
| PC 15:0_18:1 [D7] | 753.6151 | 184.0734 |
| PC 16:0_16:0 | 734.5704 | 184.0734 |
| PC 17:0_16:1 [D5] | 751.6007 | 184.0734 |
| PC 16:0_18:1 | 760.586 | 184.0734 |
| PC 17:0_18:1 [D5] | 779.632076 | 184.0734 |
| PC 18:0_18:1 | 788.6172 | 184.0734 |
| **Phosphatidylethanolamine (PE)** | | |
| PE 15:0_18:1 [D7] | 711.567 | 570.6 |
| PE 16:0_18:1 | 718.5397 | 577.5 |
| PE 17:0_18:1 [D5] | 737.5851 | 596.6 |

| | | |
|---|---|---|
| PE 18:0_18:1 | 746.5696 | 605.6 |
| **Phosphatidylserine (PS)** | | |
| PS 16:0_18:1 | 762.528 | 577.5178 |
| PS 17:0_18:1 [D5] | 803.5569 | 618.5518 |
| PS 18:0_18:1 | 790.5593 | 605.549 |
